# Structure-guided design and optimization of covalent VHL-targeted sulfonyl fluoride PROTACs

**DOI:** 10.1101/2023.06.20.545773

**Authors:** Rishi R. Shah, Elena De Vita, Daniel Conole, Preethi S. Sathyamurthi, Xinyue Zhang, Elliot Fellows, Carlos M. Fleites, Markus A. Queisser, John D. Harling, Edward W. Tate

## Abstract

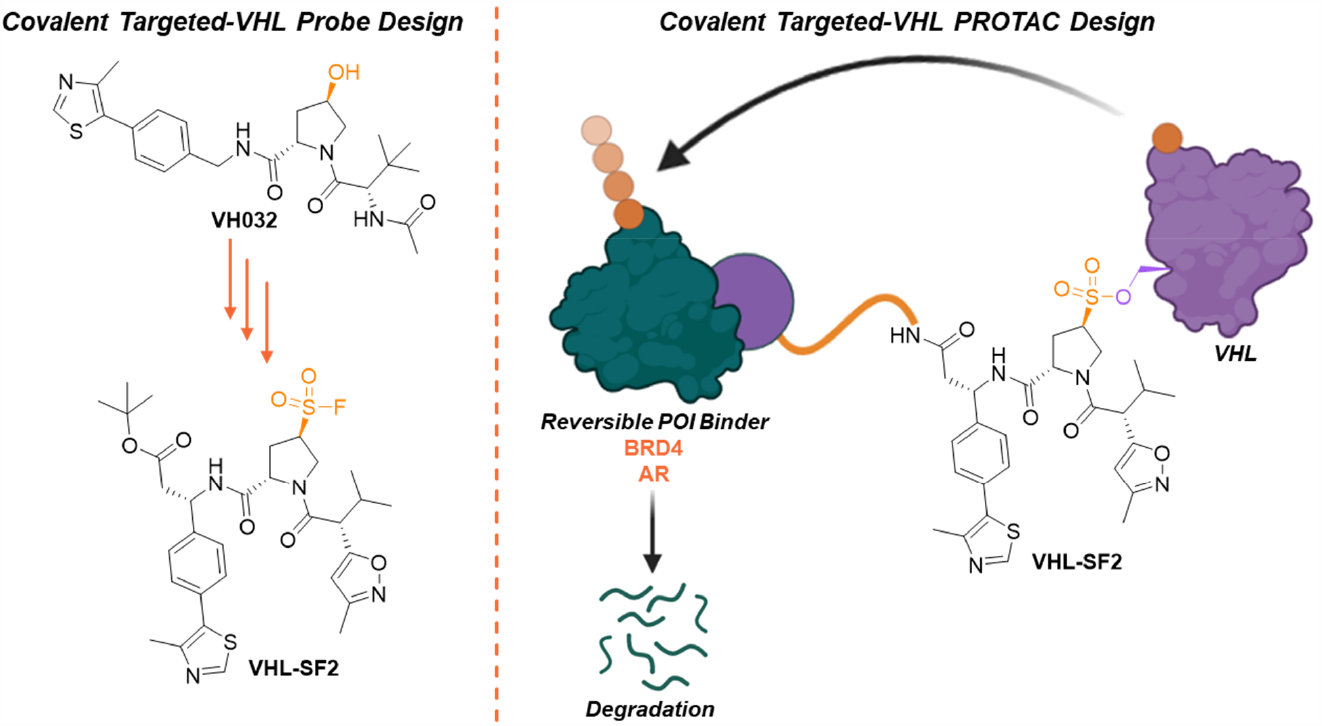

Proteolysis-targeting chimeras (PROTACs) are heterobifunctional molecules that have emerged as a therapeutic modality to induce targeted protein degradation (TPD) by harnessing cellular proteolytic degradation machinery. PROTACs which ligand the E3 ligase in a covalent manner have attracted intense interest, however, covalent PROTACs with a broad protein of interest (POI) scope have proven challenging to discover by design. Here, we report structure-guided design and optimization of Von Hippel-Lindau (VHL) protein-targeted sulfonyl fluorides which covalently bind Ser110 in the HIF1α binding site. We demonstrate that their incorporation in bifunctional degraders induces targeted protein degradation of BRD4 or androgen receptor (AR) without further linker optimization. Our study discloses the first covalent VHL ligands which can be implemented directly in bifunctional degrader design expanding the substrate scope of covalent E3 ligase PROTACs.

## Introduction

Von Hippel-Lindau (VHL) protein is among the most widely recruited E3 ligases in the PROTAC field. All potent small molecule VHL binders reported to date feature a (*R*)-hydroxyproline motif,^1,2,3,4^ which forms an essential interaction with Ser110 in the HIF1α binding site of VHL but limits passive transport across the cell membrane.^5,6,7,8^ This motif is considered essential for VHL recognition and presents a challenge for optimization of VHL PROTAC potency and cell uptake.

PROTACs which ligand the E3 ligase in a covalent manner offer potential advantages over a reversible counterpart by transforming the ternary complex into a simple binary interaction between modified E3 and substrate (Fig. 1A).^9^ To date, covalent PROTACs have been reported for DCAF1, DCAF11, DCAF16, RNF4, RNF114, and FEM1B; each bearing a cysteine-targeting electrophilic warhead (e.g. chloroacetamide or α,β-unsaturated carbonyl) and has been discovered by screening rather than by design.^10,11,12,13,14,15^ Recently reported covalent CRBN E3 ligase binders bearing a sulfonyl fluoride show intriguing molecular glue activity, although they have yet to be incorporated in a covalent PROTAC.^16^

**Figure 1.**
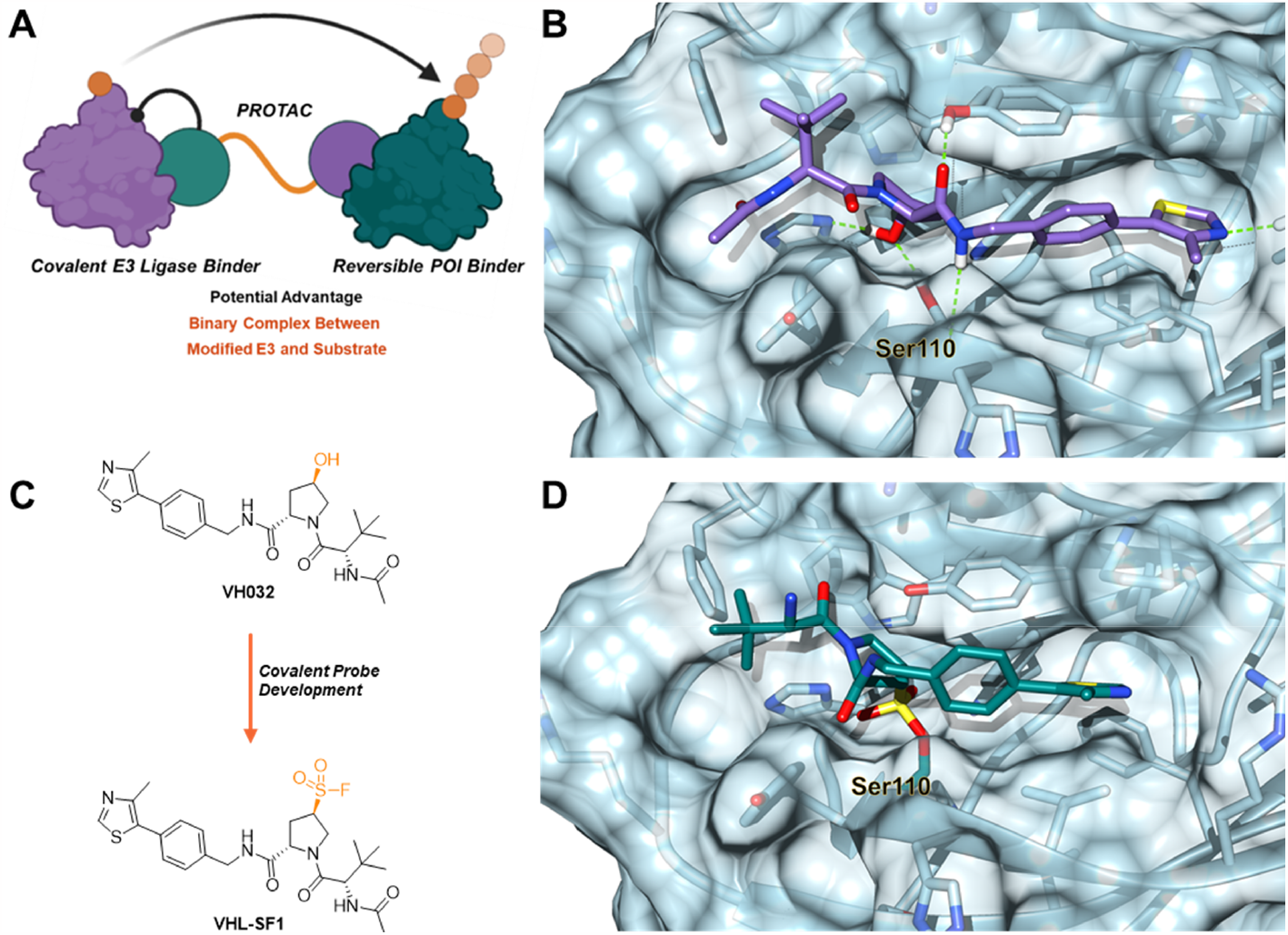
(A) Potential advantage of covalent PROTACs allowing transformation of the ternary complex into a simple binary interaction between modified E3 and substrate. (B) VH032/VHL co-crystal structure, illustrating the critical Ser110 interaction between **VH032** (purple) and VHL (PDB: 4W9H). (C) Structure of **VH032** and replacement of the hydroxyproline moiety with a sulfonyl fluoride to generate **VHL-SF1**. (D) Docking of **VHL-SF1** (blue) within a VHL crystal structure (PDB: 4W9H).

Herein, we document design and optimization of the first rationally designed covalent VHL ligands and their incorporation in PROTACs for TPD applications. We demonstrate that the hydroxyproline motif of a known VHL binder can be replaced by a sulfonyl fluoride moiety and through structure-guided optimization generate a ligand which covalently modifies Ser110 of VHL in the HIF1α binding site. We systematically assess VHL occupancy in recombinant protein and live cells, and the capacity of covalent PROTACs derived from this ligand to induce degradation of BRD4 and androgen receptor (AR). We suggest that this novel covalent VHL PROTAC paradigm will prove valuable for future studies of target engagement and optimization of cellular uptake of VHL-recruiting PROTACs.

## Results and Discussion

### First-generation sulfonyl fluoride covalent VHL ligand

The binding mode of **VH032**, the most widely exploited VHL ligand in PROTACs to date, features a critical hydrogen bond to Ser110 through the (*R*)-hydroxyproline motif (Fig. 1B).^8^ In our initial ligand designs, we sought to replace the hydroxy group with a sulfonyl fluoride, an electrophilic warhead featuring balanced reactivity, resistance to hydrolysis under physiological conditions, and the capacity to covalently modify proteins at varied nucleophilic residues beyond cysteine, including serine.^17,18^ We investigated how the hydroxyl replacement would perturb VHL binding by docking prototype covalent ligand **VHL-SF1** (Fig. 1C) in VHL derived from a VH032/VHL complex (PDB: 4W9H) (Fig. 1D, S1).^19^ These models suggest that **VHL-SF1** has the potential to covalently bind Ser110, thereby maintaining some of the critical interactions observed in the hydroxyproline motif despite a degree of displacement of the remainder of the molecule (Fig. S1), which may compromise non-covalent interactions exhibited by **VH032**. We considered **VHL-SF1** a reasonable starting point to probe covalent VHL modification.

Synthesis of **VHL-SF1** commenced with generation of (*S*)-hydroxyproline through a short sequence of reactions (Scheme S1). Displacement of mesylate **1** with thioacetate afforded compound **2**, which was converted to sulfonyl fluoride **4** via sulfonyl chloride **3** (Scheme 1). Whilst this reaction generated the desired sulfonyl fluoride, epimerization of the proline ring occurred, resulting in a mixture of diastereomers. To circumvent this, we developed novel reaction conditions to synthesize the desired sulfonyl fluoride in one step directly from thioacetate **2**, resulting exclusively in the desired stereoisomer **5**. A proposed mechanism for the novel sulfonyl fluoride transformation based on known analogous reactions is shown in Scheme S2.^20^ Following Boc-deprotection, the amine was acetylated to afford **VHL-SF1**, or biotinylated to generate **VHL-SF1-Biotin**.

To assess the ability of **VHL-SF1** to covalently modify VHL we initially developed a streptavidin shift assay in which we exposed recombinant human VCB, a stable complex of VHL protein with elongin C and elongin B, to **VHL-SF1-Biotin**. VHL biotinylation could then be directly quantified by the apparent shift in molecular weight observed when the sample was mixed with streptavidin and analyzed by anti-VHL Western blot (Fig. S2).^21^ Through this assay, we concluded that 10 μM **VHL-SF1-Biotin** modified 32% VHL following 2 hours incubation at room temperature, which was further confirmed to be concentration-dependent with respect to the probe (Fig. S3).

**Scheme 1.**
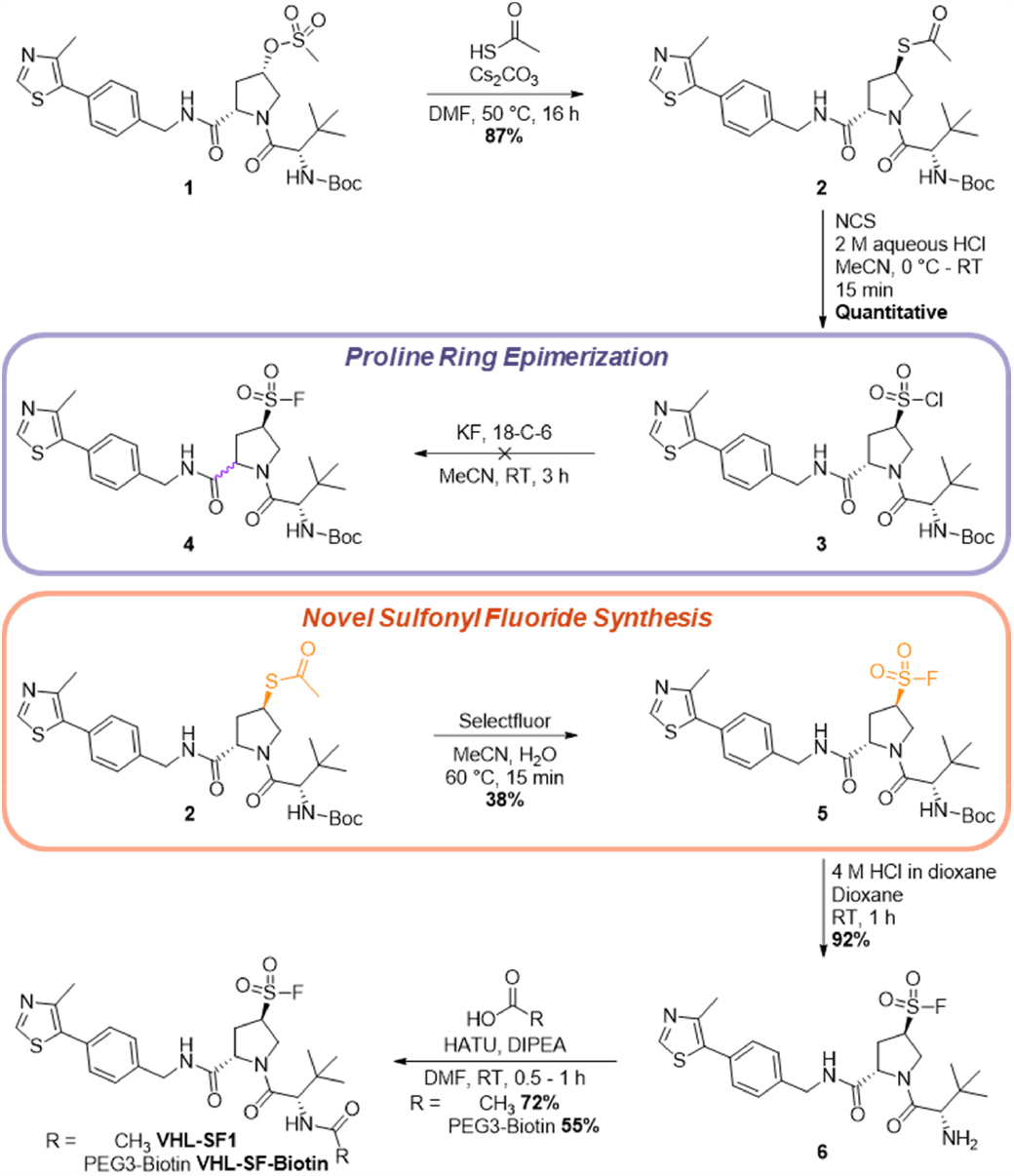
Development of a novel synthetic route for the introduction of the sulfonyl fluoride moiety in **VHL-SF1** and **VHL-SF1-Biotin**.

Consistent with this modest reactivity, **VHL-SF1** at concentrations up to 100 μM was unable to displace a fluorophore-labeled HIF1α peptide, known to occupy both the **VH032** binding site and a second VHL domain, in a competitive fluorescence polarization (FP) assay (Fig. S4). We next focused on developing a second generation covalent VHL binder with enhanced potency and occupancy.

### Second-generation sulfonyl fluoride VHL covalent ligand

In order to improve covalent ligand potency, we reasoned that optimization of the groups peripheral to the hydroxyproline motif could provide enhanced affinity for VHL. Drawing inspiration from structure-activity studies of previously reported VHL binders,^4^ we examined docked poses for analogues of **VHL-SF1**, including **VHL-SF2** in which the *tert*-leucine moiety was swapped for a methyl isoxazole, and the ligation vector moved to the benzylic position (Fig. 2A). In contrast to **VHL-SF1**, this analysis suggested that **VHL-SF2** may covalently bind Ser110 whilst maintaining many of the non-covalent interactions seen with **VH032** (Fig. 2B). **VHL-SF2** was synthesized in 11 steps (Scheme S3).

**Figure 2.**
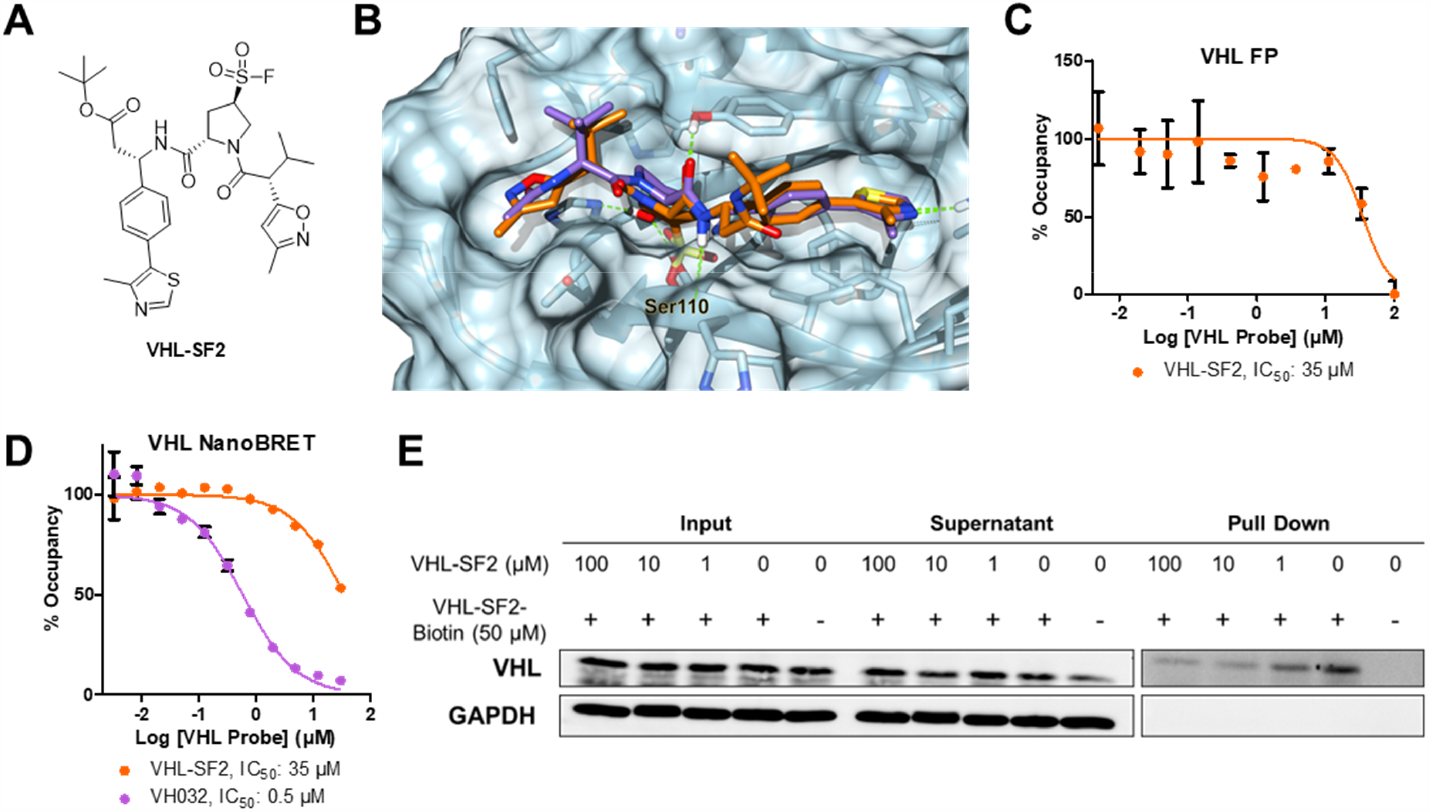
(A) Structure of **VHL-SF2**. (B) Docking of **VHL-SF2** (orange) within a crystal structure of VHL (PDB: 4W9H) superimposed onto **VH032** (purple). (C) Dose–response of **VHL-SF2** inhibition of the VCB and FAM-conjugated HIF1α-derived peptide interaction assessed by FP, following 2 h incubation between VCB and **VHL-SF2**. Data shows mean ± SEM (n = 3). (D) Cellular potency of **VHL-SF2** and **VH032** measured by a NanoBRET target engagement assay, following 5 min incubation between VHL-NanoLuc HEK293 cells, **VHL-SF2**, NanoBRET VHL tracer ligand and digitonin. Data shows mean ± SEM (n = 3). (E) Competition pull down assay from live cells between **VHL-SF2-Biotin**, and **VHL-SF2**. HEK293T cells were pre-treated with DMSO or varying concentrations of **VHL-SF2** for 2 h at 37 °C. The cells were lysed and treated with DMSO, or **VHL-SF2-Biotin** (50 μM) for 2 h at room temperature. A pull down with streptavidin beads was conducted after which protein was resolved on SDS/PAGE. VHL and loading control GAPDH levels were visualized by Western blotting.

Initially, we subjected **VHL-SF2-Biotin** to the streptavidin shift assay and observed a greater extent of VHL modification (44%) than for **VHL-SF1-Biotin** under the same conditions (Fig. S2). Furthermore, **VHL-SF2** was able to displace labeled peptide in the FP assay, with an apparent IC_50_ of 35 μM at 2 h, consistent with predicted covalent occupancy of the VHL HIF1α binding site (Fig. 2C). Intact LC-MS analysis confirmed single labeling of VHL by **VHL-SF2**, with conversion of 65% at 24 h (Fig. S5).

Encouraged by this evidence for biochemical engagement of VHL, we explored engagement of VHL by **VHL-SF2** in live cells through a competition pull-down assay from HEK293T cells (Fig. 2E). HEK293T cells were incubated with varying concentrations of **VHL-SF2** for 2 h at 37 °C, lysed and treated with **VHL-SF2-Biotin** at 50 μM for 2 h followed by pull-down on streptavidin beads. Western blot analysis showed dose-dependent competition of VHL engagement by **VHL-SF2**, consistent with covalent modification of VHL in intact cells.

Target engagement and cellular VHL binding potency were further confirmed through a NanoBRET target engagement assay, assessing inhibition of the interaction between VHL-NanoLuc and a cell-permeable fluorescent VHL tracer ligand, in HEK293 cells (Fig. 2D).^22^ In agreement with the FP assay, **VHL-SF2** inhibited the BRET signal with an IC_50_ of 35 μM, compared to >100 μM for **VHL-SF1** and 0.5 μM for **VH032**, consistent with intracellular HIF1α binding site occupancy.

### VHL-SF2-based PROTAC induces proteasome-dependent TPD of BRD4

After validating VHL engagement by **VHL-SF2** both on isolated protein and in intact cells, we synthesized a BRD4-targeted PROTAC, **BRD-SF2**, incorporating **VHL-SF2** and a known BRD4 ligand (Fig. 3A),^23^ and assessed target protein degradation using a BRD4 HiBiT assay.^22^ Pleasingly, this unoptimized PROTAC induced BRD4 degradation (DC_50_ : 17.2 μM, D_max_: 60% at 18 h incubation), whilst **VHL-SF2** alone did not affect BRD4 levels (Fig. 3B); **BRD-SF2** and **VHL-SF2** showed no evidence of cytotoxicity under these conditions (CellTiter-Glo® assay, Fig. S6A). Consistent with the BRD4 HiBiT assay, **BRD-SF2** also induced endogenous BRD4 degradation to a similar extent across the concentrations tested (BRD4 short isoform D_max_: 67%, Fig. 3C).

**Figure 3.**
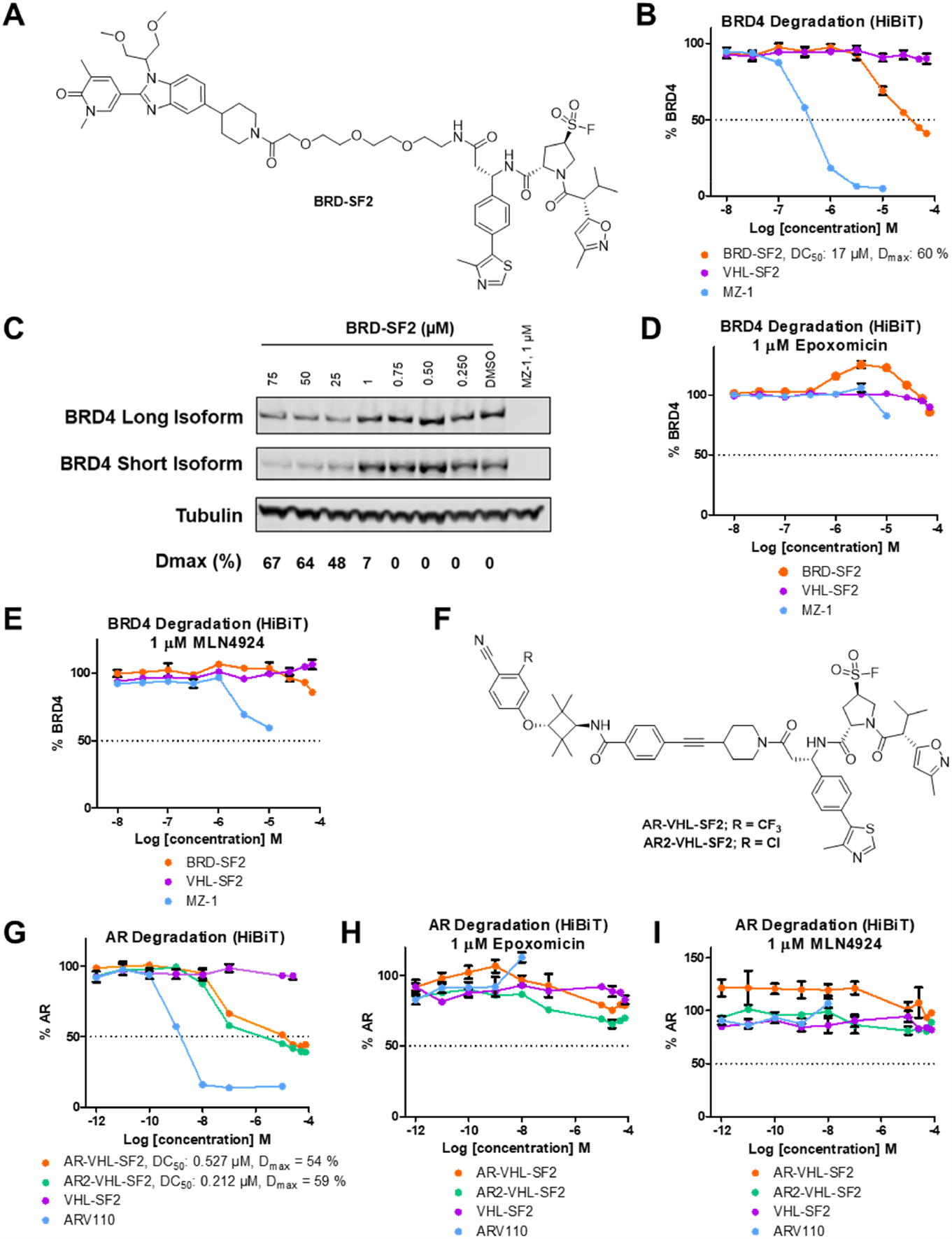
(A) Structure of **BRD-SF2**. (B) BRD4-HiBiT HEK293T cells were treated with the **BRD-SF2, MZ-1** or **VHL-SF2** dose-dependently (10 - 0.001 μM) for 18 h. Data shows representative assay from n = 4. (C) **BRD-SF2**-mediated degradation of endogenous BRD4. HEK293T cells were treated with DMSO, **MZ-1** or varying concentrations of **BRD-SF2** for 18 h. BRD4 and loading control GAPDH levels were visualized by Western blotting. Data shows representative Western blot from n = 3. (D) BRD4-HiBiT HEK293T cells were treated with epoxomicin (1 μM) for 3 h, followed by the respective PROTAC or VHL-SF2 dose-dependently (10 - 0.001 μM) for 18 h. Data shows mean ± SEM (n = 4). (E) BRD4-HiBiT HEK293T cells were treated MLN4924 (1 μM) for 3 h, followed by the respective PROTAC or **VHL-SF2** dose-dependently (10 - 0.001 μM) for 18 h. Data shows mean ± SEM (n = 4). (F) Structure of **AR-VHL-SF2** and **AR2-VHL-SF2**. (G) AR-HiBiT LNCaP cells were treated with either **AR-VHL-SF2, AR2-VHL-SF2, ARV110** or **VHL-SF2** dose-dependently (10 - 0.001 μM) for 16 h. Data shows mean ± SEM (n = 4). (H) AR-HiBiT LNCaP cells were treated with epoxomicin (1 μM) for 3 h, followed by either **AR-VHL-SF2, AR2-VHL-SF2, ARV110** or **VHL-SF2** dose-dependently (10 - 0.001 μM) for 16 h. Data shows mean ± SEM (n = 3). (I) AR-HiBiT LNCaP cells were treated with MLN4924 (1 μM) for 3 h, followed by either **AR-VHL-SF2, AR2-VHL-SF2, ARV110** or **VHL-SF2** dose-dependently (10 - 0.001 μM) for 18 h. Data shows mean ± SEM (n = 3).

BRD4 was not depleted in the presence of **BRD-SF2** when treated with inhibitors of either proteasome activity or NEDDylation (epoxomicin and MLN4924, respectively), supporting a proteasome- and Cullin E3 ligase-dependent mechanism consistent with recruitment of VHL (Fig. 3D & E).

### VHL-SF2-based PROTACs induce TPD of AR

To further confirm the versatility of **VHL-SF2**s ability to recruit VHL to induce target protein degradation when incorporated into a PROTAC, we synthesized two AR ligand-derived **VHL-SF2** conjugates, **AR-VHL-SF2** and **AR2-VHL-SF2**, based on known AR ligands with attractive cellular potency that have previously been incorporated into AR-bifunctional degraders (Fig. 3F).^24,25^ These compounds were assessed for induced AR degradation using an AR HiBiT assay in LNCaP prostate cancer cells, in which endogenous AR was tagged with a HiBiT peptide (Fig. 3G). Both **AR-VHL-SF2** and **AR2-VHL-SF2** induced AR degradation (**AR-VHL-SF2** DC_50_ = 0.527 μM, D_max_ = 54%; **AR2-VHL-SF2** DC_50_ = 0.212 μM, D_max_ = 59%), in addition to Arvinas’s AR-degrading clinical candidate, **AR110**, which was used as a positive control. Both **AR-VHL-SF2**- and **AR2-VHL-SF2**-mediated protein degradation were proteasome- and E3 ligase-dependent based on AR degradation blockade in the presence of either epoxomicin or MLN4924 (Fig. 3H & I), and did not exhibit cytotoxicity under these conditions (CellTiter-Glo® assay, Fig. S6B).

## Conclusions

To our knowledge, this is the first report of a VHL ligand that lacks the hydroxyproline motif and engages covalently with VHL via a sulfonyl fluoride moiety, and the first covalent E3 ligase binder developed by design, rather than screening. When incorporated into bifunctional degraders, the resulting sulfonyl fluoride-based PROTACs are capable of inducing proteasome- and ubiquitin ligase-dependent TPD of both BRD4 and AR without medicinal chemistry optimization. This result potentially expands the substrate scope of E3 ligase covalent PROTACs to the >20 target proteins previously reported to be degradable with VHL-recruiting PROTACs. Our work paves the way for additional structural modifications at the hydroxyproline center which could ultimately result in PROTACs targeting VHL with improved pharmacokinetic properties. This work benchmarks the covalent recruitment of VHL for TPD and provides the basis for medicinal chemistry campaigns to further optimize the covalent VHL ligand to enhance potency. We also report novel bifunctional degraders of BRD4 and AR which could be used as starting points for the development of PROTACs with improved pharmacokinetic and pharmacodynamic properties.

## Acknowledgements

The authors would like to thank Sandra Kümper, Kwok-Ho Chan, Justyna Macina, Jacob Bush, Katherine Jones, Emmanuel Demont, Peter Stacey, Eleanor Dickinson and Darren Poole for helpful discussions. We thank Chun-wa Chung for the kind donation of VCB protein and Stephen Richards for assistance with NMR. The research was supported by an EPSRC Knowledge Transfer Secondment grant by UKRI (grant C29637/A20781 to RRS and EWT). EDV is supported by a H2020 (EC) MSCA-IF, project RabTarget4Metastasis (EU project 890900). XZ is supported by a scholarship from Imperial College London and the China Scholarship Council program. Molecular graphics and analyses performed with UCSF Chimera, developed by the Resource for Biocomputing, Visualization, and Informatics at the University of California, San Francisco, with support from NIH P41-GM103311. Graphics were created with BioRender.com.

## Supplementary Information

## Supplementary Figures

**Figure S1.**
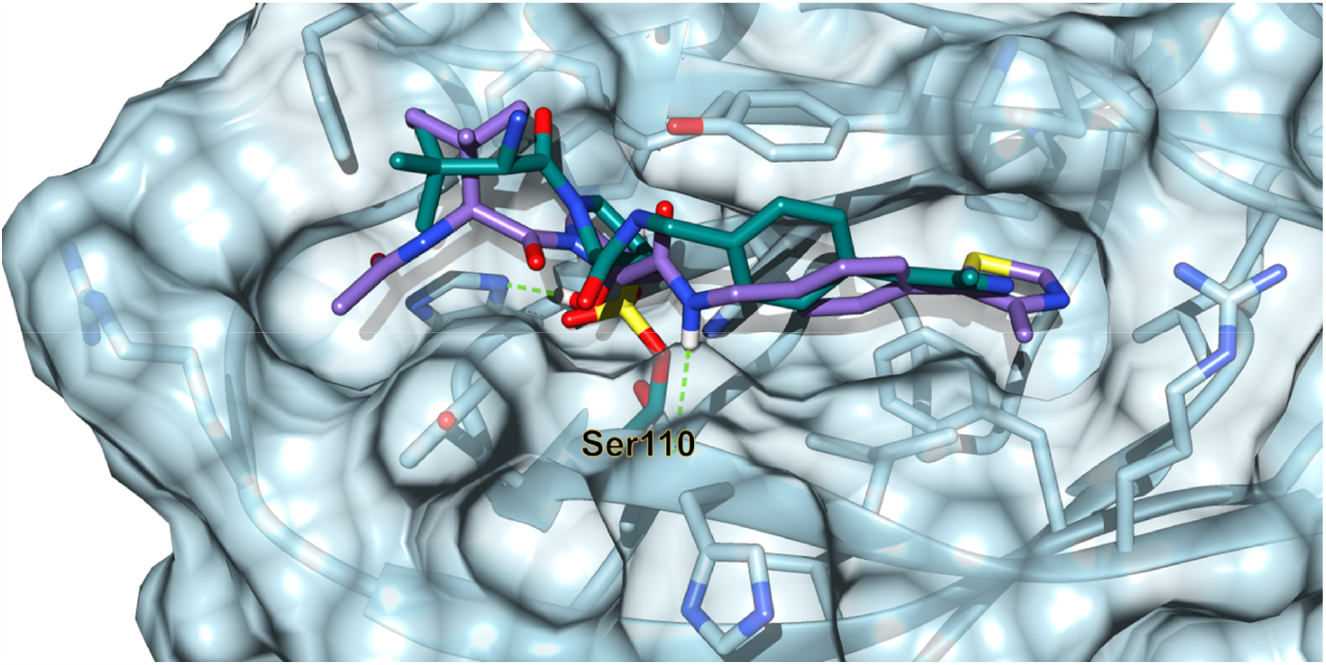
A docking of **VHL-SF1** (blue) within VHL superimposed onto **VH032** (purple). Docking was performed with MOE using the Covalent Docking protocol. Docking made use of PDB: 4W9H as the crystal structure.

**Scheme S1.**
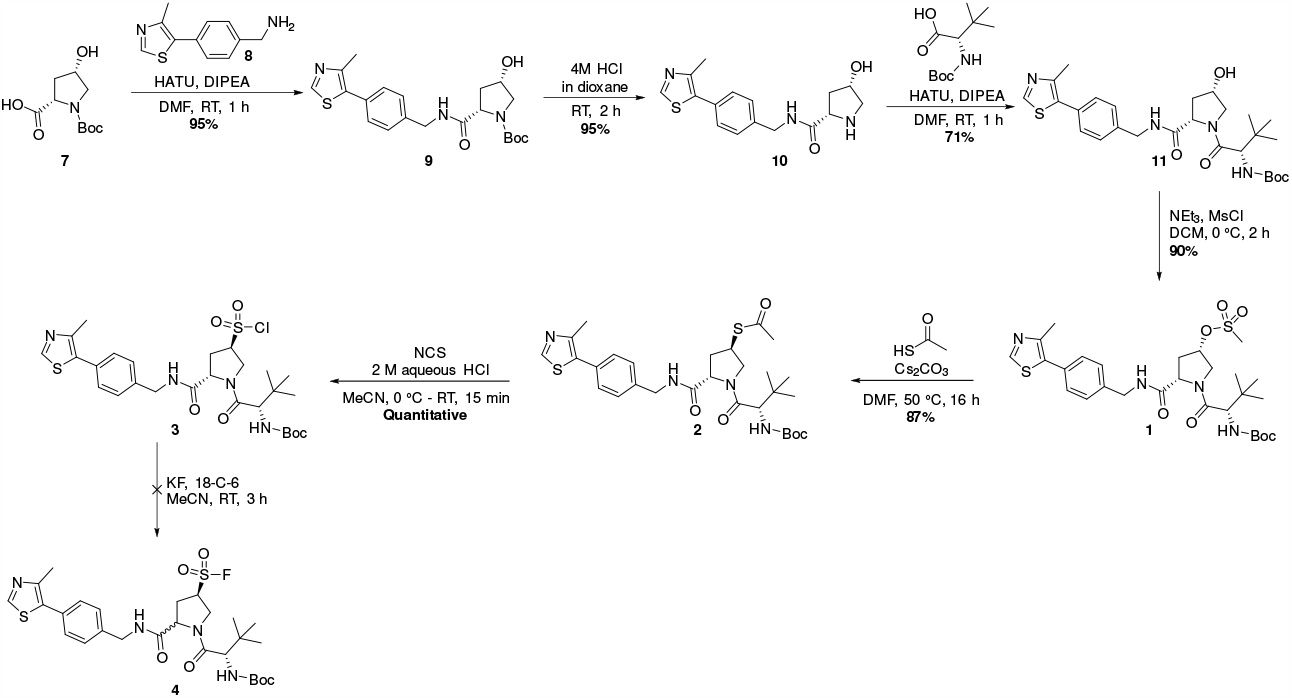
Synthesis of **VHL-SF1** resulting in epimerization of the proline ring when subjected to KF and 18-C-6.

**Scheme S2.**
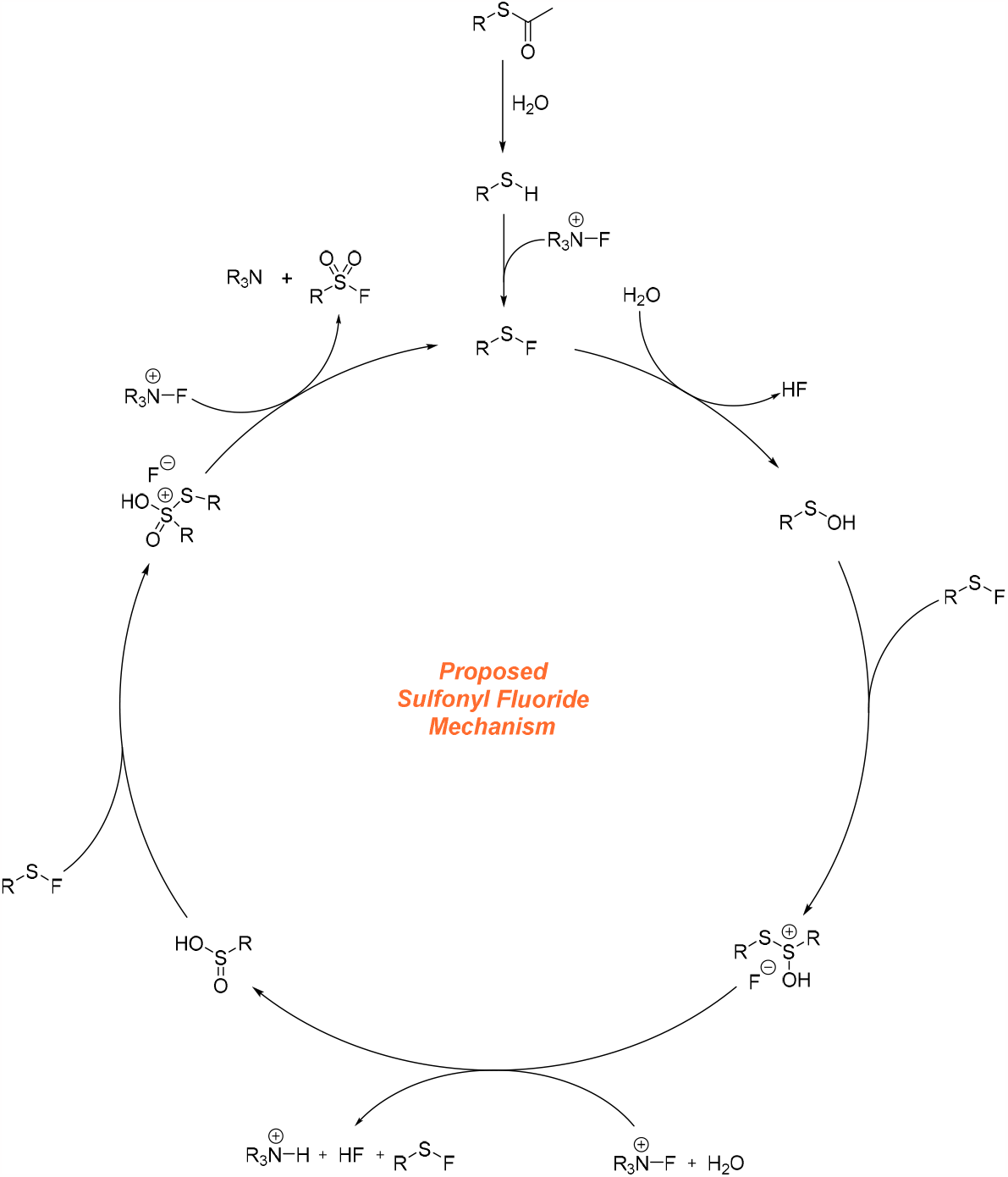
Proposed mechanism of the novel sulfonyl fluoride reaction, to convert thioacetates to sulfonyl fluorides.

**Figure S2.**
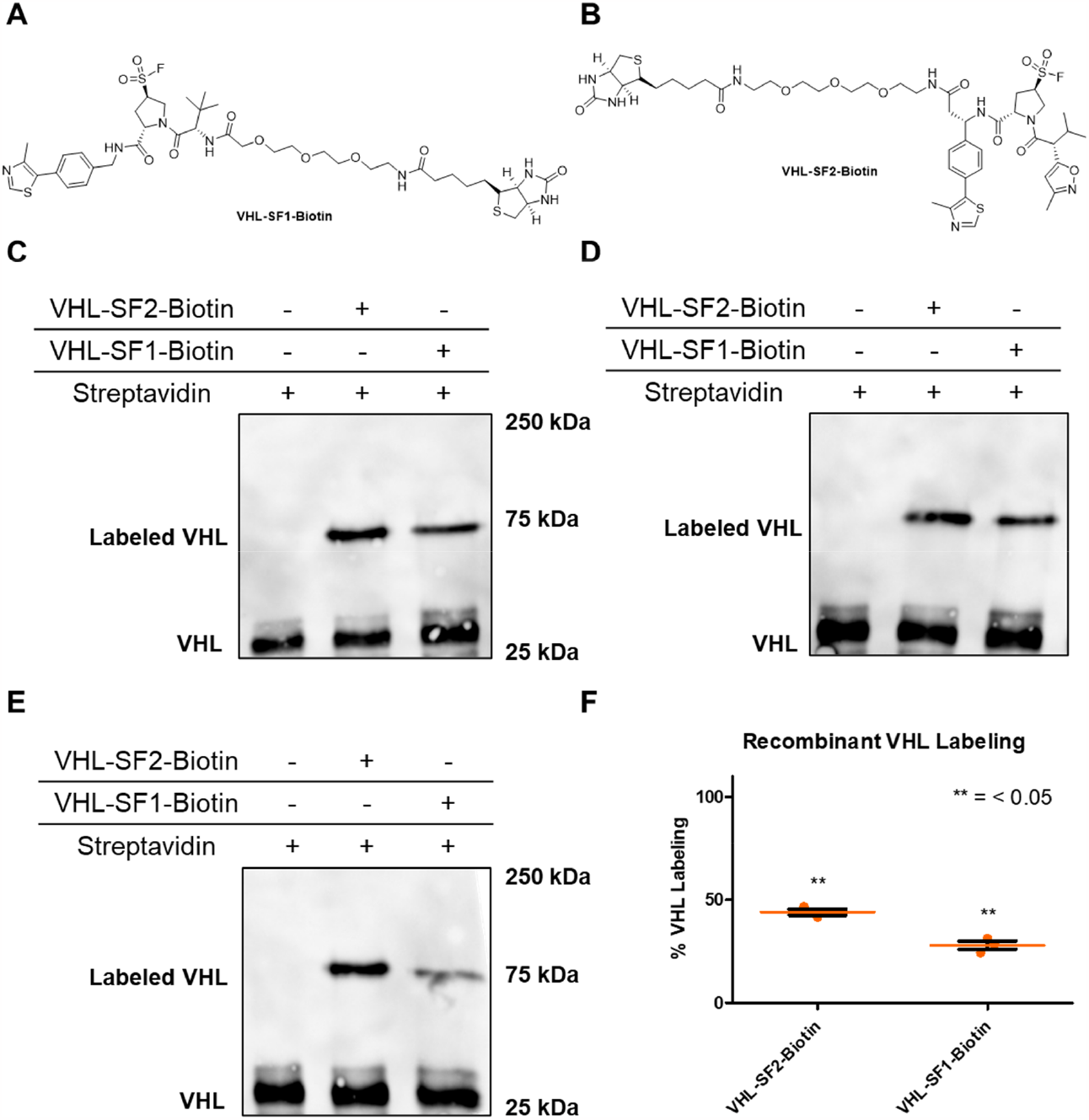
(A) Structure of **VHL-SF-1-Biotin**. (B) Structure of **VHL-SF-2-Biotin**. (C - E) Gel-based streptavidin shift assay of **VHL-SF1-Biotin**, and **VHL-SF2-Biotin**. VCB protein was pre-treated with DMSO, **VHL-SF1-Biotin** (10 μM) or VHL-SF2-Biotin (10 μM) for 2 h at room temperature prior to addition of streptavidin (10 μM, 10 min) at room temperature, after which protein was resolved on SDS-PAGE and visualised by VHL Western blotting. (F) Quantification of VHL levels from Western blot. Data shows individual quantified Western blot ± SEM (n = 3).

**Figure S3.**
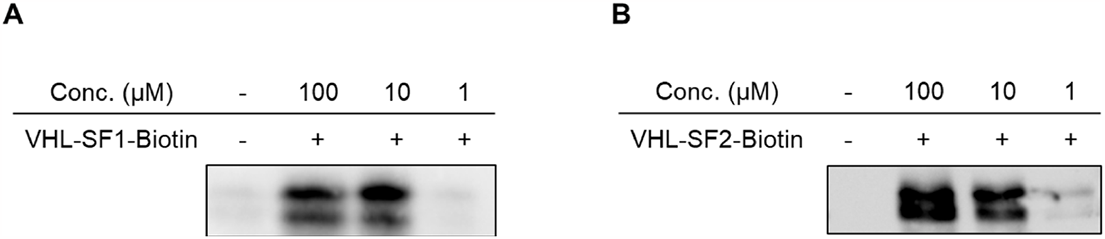
(A) Gel-based VHL labelling of **VHL-SF1-Biotin**, and (B) **VHL-SF2-Biotin**. VCB protein was pre-treated with DMSO, **VHL-SF1-Biotin** or **VHL-SF2-Biotin** for 2 h at room temperature, after which protein was resolved on SDS-PAGE and visualised by Neutravidin HRP Western blotting.

**Scheme S3.**
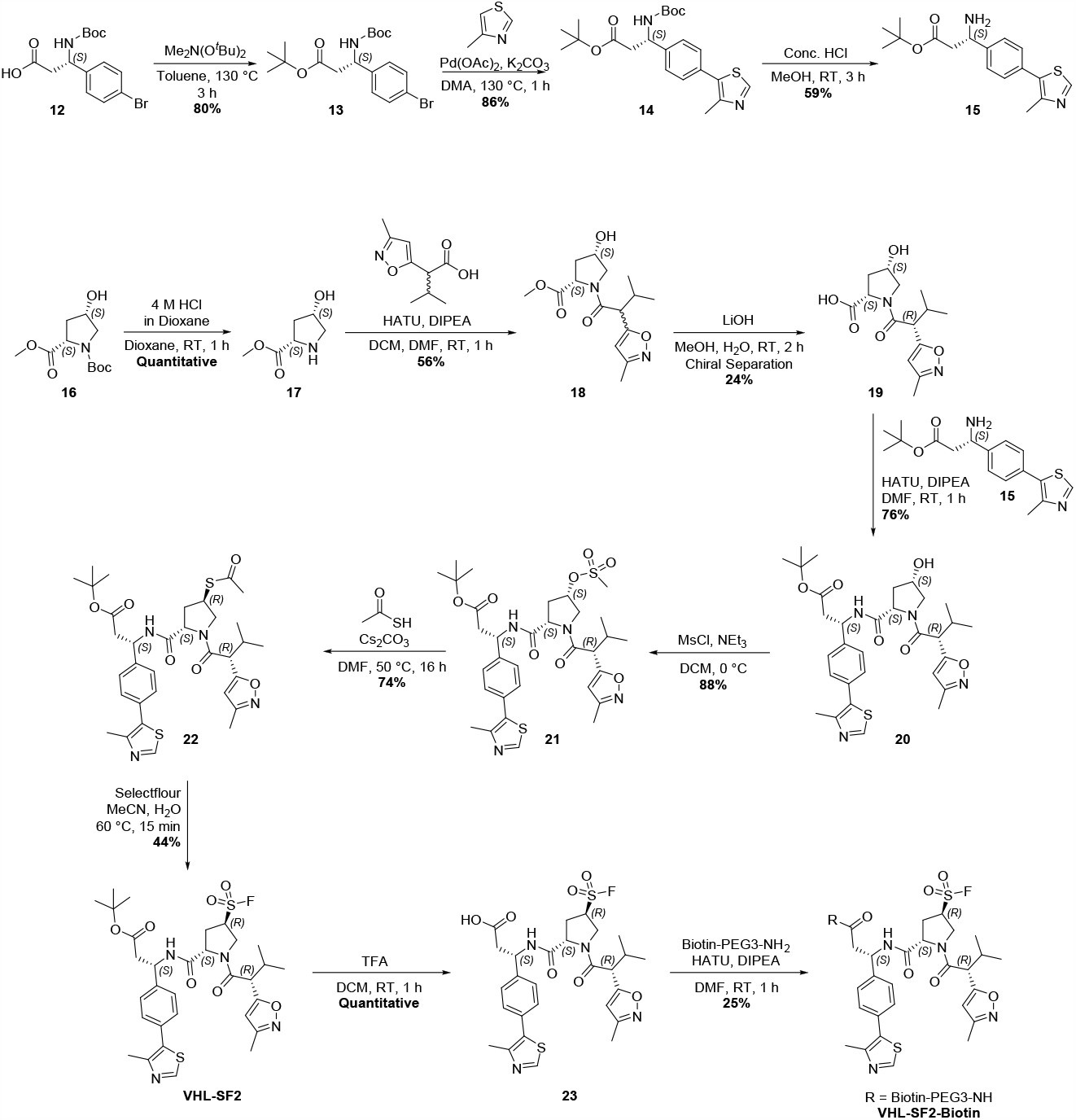
Synthesis of **VHL-SF2** and **VHL-SF2-Biotin**.

**Figure S4.**
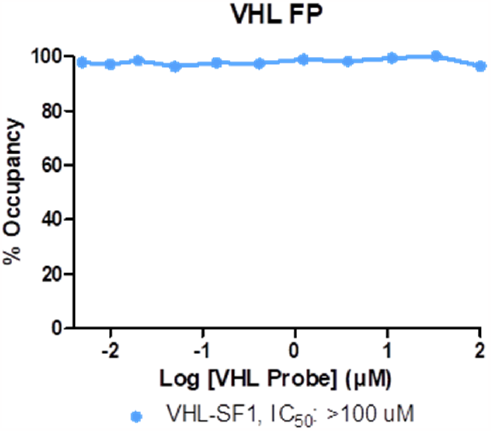
Dose-response of **VHL-SF1** following a 2 h pre-incubation in the presence of VCB and FAM-conjugated HIF1α-derived peptide, assessed by fluorescence polarization. Data shows mean ± SEM (n = 3).

**Figure S5.**
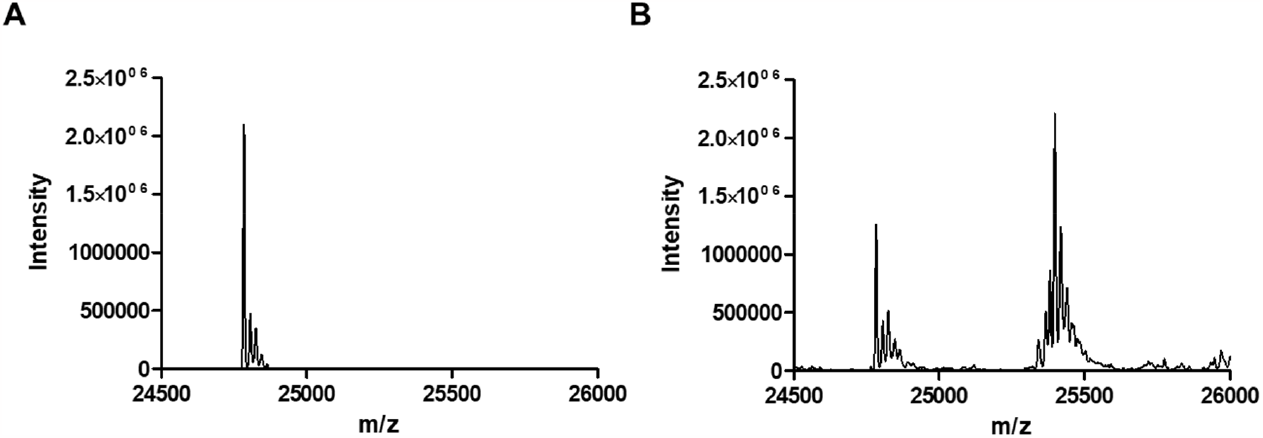
(A) VCB was incubated with DMSO for 24 h and subjected to intact-protein LC-MS (VHL: 24784 Da). (B) VCB was incubated with **VHL-SF2** (100 μM) for 24 h and subjected to intact-protein LC-MS (65% mono-VHL labelling by **VHL-SF2**; VHL: 24783, [M-^*t*^Bu+VHL+Na]^+^ 25397 Da).

**Figure S6.**
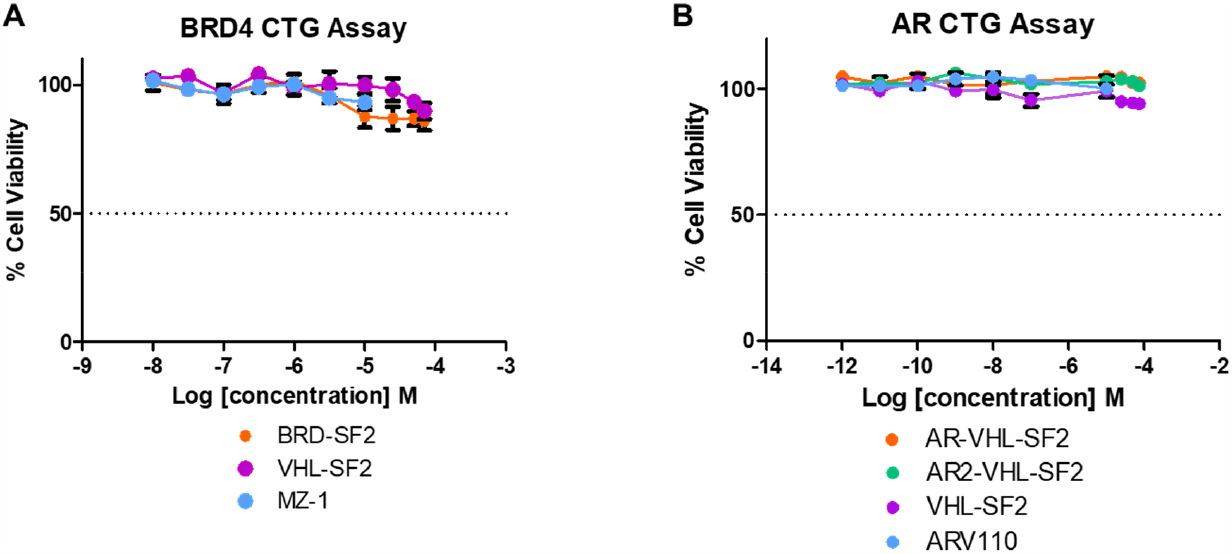
(A) BRD4-HiBiT HEK293T cells were treated with **BRD-SF2, MZ-1** or **VHL-SF2** at varying concentrations (10 - 0.001 μM) for 18 h. After the treatment period, The CellTiter-Glo^®^ 2.0 Assay reagents were used for this experiment per the manufacturer’s instruction. Data shows mean ± SEM (n = 4). (B) AR-HiBiT LNCaP Cells were treated with **AR-VHL-SF2, AR2-VHL-SF2, ARV110** or **VHL-SF2** at varying concentrations (10 - 0.001 μM) for 16 h. After the treatment period, The CellTiter-Glo^®^ 2.0 Assay reagents were used for this experiment per the manufacturer’s instruction. Data shows mean ± SEM (n = 4).

## Biological Methods

### Docking analysis with Molecular Operating Environment (MOE)

Molecular modelling studies were conducted using the software Molecular Operating Environment 2019.01. For VHL (PDB: 4W9H) docking studies, the protein was prepared using default parameters (quickprep) and the ligand site was used to perform a ‘Covalent Dock’ of VHL-SF1 and VHL-SF2, which were imported as an sdf file from ChemDraw, placing 30 poses with ‘Rigid Receptor’ and refining 5 poses with ‘GBVI/WSA dG’ scoring. The final pose was exported as a PDB file. PyMOL was used to generate docking figures.

### VHL/EloB/EloC expression and purification

A construct containing the sequence for human VHL (UniPort entry P40337, residues 1-213) was cloned into a pET151/D-TOPO vector (Invitrogen), including a *N*-terminal 6xHis-tag followed by a Tobacco Etch Virus (TEV) cleavage site. Fusion constructs, EloB and EloC (17-112), were inserted in pACYC-Duet1 vector (Addgene entry Plasmid #110274). Both plasmids were transformed into *E*.*coli* One Shot™BL21(DE3) Chemically Competent cells (Invitrogen™) and spread on LB agar plates containing 100 mg/L Ampicillin and 25 mg/L Chloramphenicol for selection. A single colony was picked and VHL/EloB/EloC was expressed at 18 °C for 18 hours after induction with 500 μM IPTG at an OD_600_ of 0.6–0.7. The cultures were spun down at 4000 rpm for 30 min at 4°C, and supernatant was removed. The collected cell pellets were resuspended in 20 mM Tris, 500 mM NaCl, 10 mM imidazole, Complete Protease Inhibitor Cocktail (EDTA-free, Roche), benzonase (2 μL/50 mL), pH 8.0 and cells were lysed using a cell disruptor at 25 kpsi followed by centrifugation at 15k g for 45 min. The supernatant was loaded on a HisTrap FF (Cytiva) chromatography equilibrated with lysis buffer, washed extensively, and eluted in a gradient of 20 mM to 500 mM imidazole. Then the obtained protein solution was added TEV protease in a molar ratio of 20:1 at 4 °C overnight in the presence of 2 mM dithiothreitol (DTT). After dialysis, the obtained protein solution was loaded on the HisTrap FF column again and the flowthrough was collected and concentrated. Finally, the protein solution was loaded on a Superdex S75 16/60 (GE Healthcare) gel filtration chromatography equilibrated in 20 mM Tris, 200 mM NaCl, 2 mM TCEP, pH 7.5 to afford the final VHL/EloB/EloC product.

### Gel-based streptavidin shift assay

VCB (11.1 mg/ mL) was diluted with PBS pH 7.4 (2 μg/ mL). VHL probes were dissolved in DMSO (500 μM) and 1.44 μL of VHL probes in DMSO (10 μM final), or 1.44 μL DMSO was added to 72 μL VCB (2 μg/ mL), and the mixtures were incubated at room temperature for 2 h. 12 μL of each sample was mixed with 4× Laemmli sample loading buffer (250 mM Tris-HCl pH 6.8, 30% (v/v) glycerol, 10% (w/v) SDS, 0.05% (w/v) bromophenol blue) supplemented with 20% v/v β-mercaptoethanol (BME) and boiled for 10 min at 95 °C. 1 mg streptavidin (Jackson Immunoresearch, Cat# 016-000-084) was dissolved in water (181 μL) to obtain a stock solution of 100 μM. 1.2 μL of 100 μM streptavidin water solution (10 μM final) was added to the appropriate samples and the mixtures were incubated at room temperature for 10 min. 10 μL of each sample was loaded onto the gel and the bands were quantified using ImageJ.

### Gel-based labelling of VHL

VCB (11.1 mg/ mL) was diluted with PBS pH 7.4 (2 μg/ mL). VHL probes were dissolved in DMSO (5 mM, 500 μM and 50 μM) and 1.44 μL of VHL probes in DMSO (100 μM, 10 μM or 1 μM final), or 1.44 μL DMSO was added to 72 μL VCB (2 μg/ mL), and the mixtures were incubated at room temperature for 2 h. 12 μL of each sample was mixed with 4× Laemmli sample loading buffer (250 mM Tris-HCl pH 6.8, 30% (v/v) glycerol, 10% (w/v) SDS, 0.05% (w/v) bromophenol blue) supplemented with 20% v/v β-mercaptoethanol (BME) and boiled for 10 min at 95 °C. 10 μL of each sample was loaded onto the gel and the bands were quantified using ImageJ.

### Fluorescence polarization assay protocol

VHL probes were dissolved in DMSO (5 mM) and diluted 25-fold with VHL assay buffer (PBS pH 7.4). The compounds were then diluted 3-fold with 4% DMSO in VHL assay buffer 9 times. 0.72 μL of 220.26 μM VCB was diluted with VHL assay buffer (160 nM). FAM-DEALAHypYIPMDDDFQLRSF was dissolved in DMSO (10 μM). 1.9 μL of 10 μM FAM-DEALAHypYIPMDDDFQLRSF (DMSO) was diluted 500-fold with VHL assay buffer (20 nM). FAM-DEALAHypYIPMDDDFQLRSF was used a positive control. For polarization displacement, 5 μL of the diluted VHL probes, and 5 μL of 160 nM VCB (80 nM Final) were added to a 384 well plate (Corning™ 3575). The plate was incubated at room temperature for 2 h. 10 μL of 20 nM FAM-DEALAHypYIPMDDDFQLRSF (10 nM Final) was added to the 384 well plate and the plate was shaken for 1 min, before reading fluorescence polarization on a Perkin Elmer Envision 2101 plate reader (excitation 486 nM, emission 535 nM). Wells containing VCB, DMSO vehicle, FAM-DEALAHypYIPMDDDFQLRSF served as maximum polarization (or minimum displacement). Wells containing buffer in place of VCB, DMSO vehicle, FAM-DEALAHypYIPMDDDFQLRSF served as minimum polarization (or maximum displacement). The percent inhibition was determined by normalizing to maximum and minimum polarization, and graphed against the log [VHL probes]. IC_50_ values were determined using Prism 5 for each replicate (n = 3), which were then averaged to determine the average IC_50_ and the standard error of the mean (SEM).

### VHL Reactivity Mass Spectroscopy

VHL-SF2 (100 μM Final) or DMSO (2% Final) was combined with VCB (1 μM) in buffer (pH 7.5, 25 mM HEPES, 150 mM NaCl) and incubated at room temperature for 24 h. The plate was centrifuged (1000 rpm, 1 min) and then subjected to intact-protein LC–MS analysis.

### Cell culture

For the gel-based competition pull down assay HEK293T cell line were obtained from The Francis Crick Institute cell services core facility and were cultured in DMEM high glucose media, supplemented with 10% (v/v) FBS, incubated at 37 °C in a 5% CO_2_ humidified incubator. Cells were grown in 75 cm^2^ cell culture flasks and 6 well plates (Corning™ 3506) for treatment. Cells were detached with trypsin (0.25%) during passaging.

### Gel-based competition pull down assay

HEK293T cells were plated into two 6 well plates (in 1 mL media 10% FBS low glucose media) at a cell density of 1.0 × 10^6^ cells/well 24 h before treatment. The cells were treated with either 2 μL 50 mM VHL-SF2 in DMSO (100 μM final), 2 μL 5 mM VHL-SF2 in DMSO (10 μM final), 2 μL 5 μM VHL-SF2 in DMSO (1 μM final), or 2 μL DMSO. The plates were swirled and then incubated at 37 °C for 2 h. The media was removed, the cells were briefly washed twice with PBS and 0.5% NP-40 (200 μL) was added to each well. The cells were scraped and let on ice for 10 min, then the suspension was centrifugated (17,000g x 10 min at 4 °C) and the supernatant was collected. Samples were adjusted to 1 mg/ mL using a Bio-Rad *DC*™ Protein Assay (#5000112). 200 μL lysates were treated with 2 μL 5 mM VHL-SF2-Biotin in DMSO (50 μM final) and incubated at room temperature for 2 h. Cold acetone (400 uL) was added, and the samples were stored at -20 °C to allow for protein precipitation. Samples were centrifuged (17,000g x 4 min at 4 °C), the supernatant was discarded and the pellet was washed once in cold acetone before being resuspended in 0.2% SDS in PBS (220 uL). An aliquot (15 μL) was kept for total lysate input before pulldown. The samples (200 μL) were incubated with pre-washed Streptavidin Magnetic Beads (New England Biolabs #S1420S, 30 μL beads per sample) for 2 h at room temperature. An aliquot (15 μL) of the supernatant was collected. The beads were washed with 0.2% (w/v) SDS in PBS pH 8.0 (1 mL × 5) and eluted by boiling in 1× Laemmli buffer (95 °C, 10 min). Proteins were separated by SDS-PAGE and further analyzed by immunoblotting.

### VHL NanoBRET TE Assay

HEK293 cells (10 × 10^6^ cells at 2 × 10^5^ cells/ mL) were transfected with 2.6 μg of VHL-NanoLuc plasmid (PROMEGA) plus 24 μg of transfection carrier plasmid (PROMEGA) and incubated for 20 h at 37 °C, in a 5% CO_2_ incubator. Cells were trypsinized, resuspended in Opti-MEM I reduced serum medium and plated into 384-well non-binding surface plates. 0.25 μM VHL tracer (PROMEGA) was added. Compounds were titrated in 11-point dilution series from 30 μM to 3 nM. Immediately after addition of tracer and compounds, cells were treated with digitonin after which NanoBRET Nano-Glo Substrate and inhibitor were added and BRET signal was measured in a Clariostar plate reader (BMG Labtech) at 450 nm and 610 nm within 5 minutes after addition of digitonin. BRET signal versus compound concentration data were fitted to a three parameters dose response logistic model fixing top to the signal in the presence of tracer and absence of compound and the bottom to the signal in the absence of tracer and compound.

### HiBiT cell lines

HEK293T cells (Takra #632180) were maintained in DMEM, high glucose, GlutaMAX™, pyruvate (ThermoFisher # 31966021) containing 10% (v/v) Fetal Bovine Serum, qualified, heat inactivated, Australia (Gibco #10100147), Penicillin-Streptomycin (10,000 U/mL) (Gibco #15140-122). AR-HiBiT KI LNCaP Cells were purchased from Promega. The cells were maintained in RPMI-1640 (Gibco # 32404-014) containing 10% (v/v) Fetal Bovine Serum, qualified, heat inactivated, Australia (Gibco #10100147), GlutaMAX™ (Gibco #35050-038) Sodium pyruvate (Gibco #11360-039) 1XPenicillin-Streptomycin (10,000 U/mL) (Gibco #15140-122). Cells were grown at 37°C with 5% CO_2_ in a humidified incubator. For passaging, cells were incubated with TrypLE™ Express Enzyme (1X), no phenol red (Gibco #12604013) at 37 °C to detach cells. For the treatments 1 μM of Epoxomicin (Sigma #E3652-50UG) and MLN4924 (Millipore) were used. The cells were pre-treated with the Epoxomicin or MLN4924 for 3 hours before the treatment with the compounds.

### Cell viability assay

The number of viable cells in the assay was determined based on the quantitation of the ATP present, which signals the presence of metabolically active cells. HEK293T HiBiT-BRD4 Cln3 and the AR-HiBiT KI LNCaP Cells were plated 10, 000 cells per well in 384-well format. The cells were treated with indicated compounds for 18h. After the treatment period, The CellTiter-Glo® 2.0 Assay reagents were used for this experiment per the manufacturer’s instruction. The plate was incubated for 10 min on an orbital shaker (at 600 RPM), and luminescence was read using a PHERAstar FS plate reader (BMG Labtech). Corresponding background RLU values were subtracted from DMSO control values and the percentage (%) of cell viability were then calculated against DMSO-treated controls. Data was analysed using Excel (Microsoft) and Prism software (v9.4.0). The assay was performed in parallel to the Nano-Glo HiBiT lytic detection assay.

### SDS-PAGE

Typically, protein or lysate samples (10–20 μg) were mixed with 4× Laemmli sample loading buffer (250 mM Tris-HCl pH 6.8, 30% (v/v) glycerol, 10% (w/v) SDS, 0.05% (w/v) bromophenol blue) supplemented with 20% v/v β-mercaptoethanol (BME) and boiled for 10 min at 95 °C. The protein or lysate samples were separated on 12% (w/v) acrylamide TrisHCl gels using Tris/glycine/SDS running buffer (0.25 M Tris, 0.2 M glycine, 0.1% (w/v) SDS) at 90 V for 15 min followed by 150 V for 1 h. Protein MW markers: Precision Plus All Blue Standards or Precision Plus Protein Dual Color Standards, Bio-Rad.

### Immunoblotting

Gels were briefly washed with deionised water, and the proteins were then transferred to a 0.45 μm nitrocellulose membrane (Amersham™ Protran®, GE Healthcare) using a wet-tank transfer (Bio-Rad) in Tris-Glycine transfer buffer (25 mM Tris, 190 mM glycine and 20% v/v MeOH) for 1 h at 100 V. For the streptavidin shift and pull down assays membranes were blocked in 5% (w/v) dried skimmed milk in Tris-buffer (50 mM Tris pH 7.4, 150 mM NaCl) containing 0.1% (v/v) Tween-20 (TBS-T) for 1 h at room temperature before incubation with the appropriate primary antibody in 5% (w/v) dried skimmed milk in TBS-T overnight at 4 °C (Table S1). The membranes were washed three times with TBS-T for 5 min and incubated with the corresponding HRP-conjugated secondary antibody (α-rabbit-HRP) in 5% (w/v) dried skimmed milk in TBS-T for 1 h at room temperature. For the Gel-based VHL-labeling assay, membranes were blocked in 3% (w/v) BSA in Tris-buffer (50 mM Tris pH 7.4, 150 mM NaCl) containing 0.1% (v/v) Tween-20 (TBS-T) overnight at 4 °C before incubation with NeutrAvidin™, Horseradish Peroxidase conjugate, #A2664, 1:1000) in 3% (w/v) BSA in TBS-T for overnight at 4 °C. For all membranes, after washing with TBS-T (5 min, ×3), the membranes were incubated with HRP substrate (Luminata Crescendo, Millipore) and the chemiluminescence signal captured with an ImageQuant™ LAS 4000 imager.

**Table S1.** Primary and Secondary antibodies were used at indicated concentrations for immunoblotting as described in Biological Methods.

| Target | Species | Dilution | Catalog No. | Supplier |
| --- | --- | --- | --- | --- |
| VHL | Rabbit | 1:1000 | 68547 | Cell Signalling |
| GAPDH | Rabbit | 1:2500 | ab9485 | Abcam |
| Rabbit IgG, HRP | Rabbit | 1:10000 | R-05072-500 | Advansta |

### BRD4 Degradation SDS-PAGE and Western blotting

Following treatment of HEK293T cells with the appropriate PROTAC and varying concentrations, the cells were lysed in RIPA buffer (ThermoFisher #89900) supplemented with Complete protease inhibitor cocktail (Roche), PhosSTOP (Roche), Benzonase (Sigma 1:1000) and 1mM DTT (Sigma #43816). After incubation for 20 min on ice, lysates were clarified by centrifugation at 17,000 xg for 20 min at 4C. Protein concentration was determined according to the Pierce™ BCA Protein Assay to enable normalisation between samples. The Lysates were processed further for SDS-PAGE. Cell extracts (25 μg total protein) were resolved by SDS-PAGE and transferred to PVDF membranes. For Western blotting, PVDF membranes were blocked in Intercept® (TBS) Blocking buffer (Licor) for 1 h at room temperature and incubated overnight at 4 °C in Intercept® (TBS) Blocking buffer (Licor) with the appropriate primary antibodies. The primary antibodies used were Recombinant Anti-Brd4 antibody [EPR5150(2)] Abcam #ab128874 1:1000 WB), Tubulin (Sigma #4967S 1:5000 WB). Membranes were subsequently washed with TBS-T and, incubated with IRDye® 800CW Donkey anti-Rabbit IgG Secondary Antibody (Licor #926-32213) and IRDye® 680RD Donkey anti-Mouse IgG Secondary Antibody (Licor #926-68072) secondary antibody for 1 h at room temperature. After further washing, signal detection was observed using LI-COR Odyssey imaging system, and signals were quantified using Image Studio software (LI-COR).

## Synthetic Chemistry

### General Methods

All reagents were used as received from commercial sources (Sigma Aldrich, Enamine, Combi-blocks etc.), unless otherwise stated. In all syntheses, anhydrous solvents were used, and commercially available HPLC grade solvents were used for work-up and isolation procedures, unless otherwise stated.

### Nuclear Magnetic Resonance Spectroscopy (NMR)

NMR spectra were recorded using a Bruker AV-400 (^1^H = 400 MHz, ^13^C = 101 MHz, ^19^F = 376 MHz), AV-500 (^1^H = 500 MHz, ^13^C = 126 MHz) or AV-600 (^1^H = 600 MHz, ^13^C = 151 MHz). Chemical shifts (δ) are reported in parts per million (ppm) relative to tetramethylsilane, DMSO, CHCl_3_ or CH_3_OD and coupling constants (*J*) in Hz. The following abbreviations are used for multiplicities: s = singlet; br. s = broad singlet; d = doublet; t = triplet; q = quartet; app. q = apparent quartet; m = multiplet; dd = doublet of doublets; dt = doublet of triplets. If not specifically stated, the NMR experiments were run at 30 °C and ^19^F and ^13^C were run in {^1^H}-decoupled mode.

### Liquid Chromatography Mass Spectrometry (LCMS)

Final LCMS analyses were conducted using one of the three methods below.

### LCMS Method A

The liquid chromatography (LC) analysis was conducted on an Acquity UPLC CSH C_18_ column (50 mm x 2.1 mm internal diameter, 1.7 μm packing diameter) at 40 °C using a 0.5 μL injection volume.

The solvents employed were:

A = 0.1 % v/v solution of formic acid in water.

B = 0.1 % v/v solution of formic acid in acetonitrile.

The gradient employed was:

| Time (min) | Flow Rate ($\text{mL min}^{-1}$) | % A | % B |
| --- | --- | --- | --- |
| 0.00 | 1 | 97 | 3 |
| 1.50 | 1 | 5 | 95 |
| 1.90 | 1 | 5 | 95 |
| 2.00 | 1 | 97 | 3 |

The UV detection was a summed signal from a wavelength of 210 nm to 350 nm. Mass spectra were recorded on a Waters ZQ mass spectrometer using alternate-scan positive and negative electrospray ionisation (ES^+^ and ES^−^) with a scan range of 100 to 1000 amu, scan time of 0.27 s and an inter-scan delay of 0.10 s.

### LCMS Method B

The liquid chromatography (LC) analysis was conducted on an Acquity UPLC CSH C_18_ column (50 mm x 2.1 mm internal diameter, 1.7 μm packing diameter) at 40 °C using a 0.3 μL injection volume.

The solvents employed were:

A = 10 mM ammonium bicarbonate in water adjusted to pH 10 with ammonia solution.

B = Acetonitrile.

The gradient employed was:

| Time (min) | Flow Rate (mL min<sup>-1</sup>) | % A | % B |
| --- | --- | --- | --- |
| 0.00 | 1 | 97 | 3 |
| 0.05 | 1 | 97 | 3 |
| 1.50 | 1 | 5 | 95 |
| 1.90 | 1 | 5 | 95 |
| 2.00 | 1 | 97 | 3 |

The UV detection was a summed signal from a wavelength of 210 nm to 350 nm. Mass spectra were recorded on a Waters ZQ mass spectrometer using alternate-scan positive and negative electrospray ionisation (ES^+^ and ES^−^) with a scan range of 100 to 1000 amu, scan time of 0.27 s and an inter-scan delay of 0.10 s.

### High Resolution Mass Spectrometry (HRMS)

High-resolution mass spectra were recorded on a Micromass Q-ToF Ultima hybrid quadrupole time-of-flight mass spectrometer, with analytes separated on an Agilent 1100 Liquid Chromatography equipped with a Phenomenex Luna C_18_ (2) reversed phase column (100 mm x 2.1 mm, 3 μm packing diameter). LC conditions were 0.5 mL·min^−1^ flow rate, 35 °C, injection volume 2 – 5 μL. Gradient elution with (A) water containing 0.1% (v/v) formic acid and (B) acetonitrile containing 0.1% (v/v) formic acid. Gradient conditions were initially 5% B, increasing linearly to 100% B over 6 min, remaining at 100 % B for 2.5 min then decreasing linearly to 5% B over 1 min followed by an equilibration period of 2.5 min prior to the next injection. Mass to charge ratios (*m/z*) are reported in Daltons.

### Mass Directed Automated Preparative HPLC (MDAP)

MDAP purifications were conducted on a Waters FractionLynx system comprising of a Waters 600 pump with extended pump heads, Waters 2700 autosampler, Waters 996 diode array and Gilson 202 fraction collector. The high performance liquid chromatography (HPLC) separation was conducted on an Xselect C_18_ column (150 mm x 30 mm internal diameter, 5 μm packing diameter) at ambient temperature, utilising an appropriate solvent system and elution gradient as determined by analytical LCMS (i.e. formic acid or ammonium bicarbonate modifier). Mass spectra were recorded on a Waters ZQ mass spectrometer using alternate-scan positive and negative electrospray ionisation (ES^+^ and ES^−^) with a scan range of 150 to 1500 amu, scan time of 0.50 s and an inter-scan delay of 0.25 s. The software used was MassLynx 3.5 with FractionLynx 4.1.

### Column Chromatography

Automated column chromatography was conducted on a Teledyne Isco Combiflash Rf system using RediSep Rf Silica cartridges (for normal phase), or Biotage KP-C_18_-HS cartridges (for reverse phase) of the correct size. Elution utilised standard HPLC grade solvents provided by Sigma Aldrich, with the desired modifier (for reverse phase) added in-house, unless otherwise stated.

### Synthesis of VHL-SF1

#### *Tert*-butyl(2*S*,4*S*)-4-hydroxy-2-((4-(4-methylthiazol-5-yl)benzyl)carbamoyl)pyrrolidine-1-carboxylate (9)

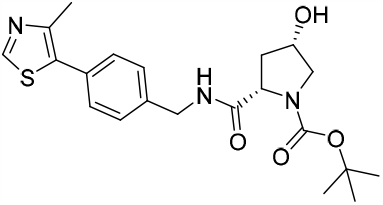

HATU (4.93 g, 12.97 mmol) was added to a stirred solution of (2*S*,4*S*)-1-(*tert*-butoxycarbonyl)-4-hydroxypyrrolidine-2-carboxylic acid (2 g, 8.65 mmol), DIPEA (3.02 mL, 17.30 mmol) and (4-(4-methylthiazol-5-yl)phenyl)methanamine (1.77 g, 8.65 mmol) in DMF (9.25 mL) and the reaction mixture was stirred at room temperature for 1 h, within a sealed vessel. The reaction mixture was diluted with water (10 mL) and extracted with DCM (3 × 10 mL). The organic layers were combined, passed through a hydrophobic frit and the solvent removed in vacuo. The residue was purified by reverse phase column chromatography (15 - 85% MeCN in H_2_O + 0.1% NH_4_HCO_3_, 330 g C18, 10 CV) to afford *tert*-butyl (2*S*,4*S*)-4-hydroxy-2-((4-(4-methylthiazol-5-yl)benzyl)carbamoyl)pyrrolidine-1-carboxylate (3.431 g, 8.22 mmol, 95% yield) as a yellow solid. ^**1**^**H NMR** (400 MHz, DMSO-*d*_6_) δ = 8.98 (1H, s), 8.50 (1H, t, *J* = 6.15 Hz), 7.37 - 7.44 (4H, m), 5.18 - 5.30 (1H, m), 4.33 - 4.43 (2H, m), 4.24 - 4.31 (1H, m), 4.08 - 4.14 (1H, m), 3.50 (1H, dd, *J* = 10.83, 5.41 Hz), 3.17 - 3.27 (1H, m), 2.44 (3H, s), 2.30 - 2.37 (1H, m), 1.73 - 1.83 (1H, m), 1.41 (3H, s), 1.26 (6H, s); ^**13**^**C NMR** (151 MHz, DMSO-*d*_6_) δ = 173.4, 153.8, 152.0, 151.9, 148.3, 139.8, 129.3 (2C), 129.2, 128.6 (2C), 127.9, 79.2, 68.5, 59.5, 54.9, 42.4, 39.2, 38.2, 28.4, 16.3; **LCMS** (Method B): t_R_ = 0.88 min, [M+H]^+^ 418, (93% purity); **HRMS** (C_21_H_28_N_3_O_4_S) [M+H]^+^ requires 418.1801, found [M+H]^+^ 418.1805.

#### (2*S*,4*S*)-4-Hydroxy-*N*-(4-(4-methylthiazol-5-yl)benzyl)pyrrolidine-2-carboxamide, 2HCl (10)

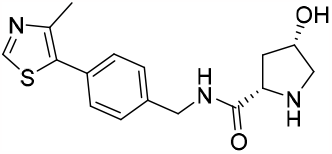

4 M HCl in Dioxane (50.7 mL, 203 mmol) was added to *tert*-butyl (2*S*,4*S*)-4-hydroxy-2-((4-(4-methylthiazol-5-yl)benzyl)carbamoyl)pyrrolidine-1-carboxylate (3.388 g, 8.11 mmol) and the reaction mixture was stirred at room temperature 2 h. The solvent was removed in vacuo to afford (2*S*,4*S*)-4-hydroxy-*N*-(4-(4-methylthiazol-5-yl)benzyl)pyrrolidine-2-carboxamide, 2HCl (3.017 g, 7.73 mmol, 95% yield) as an ivory solid. ^**1**^**H NMR** (400 MHz, DMSO-*d*_6_) δ = 8.98 10.00 - 10.18 (1H, m), 9.10 (1H, t, *J* = 6.15 Hz), 9.04 (1H, s), 8.52 - 8.67 (1H, m), 7.35 - 7.51 (4H, m), 4.35 - 4.48 (4H, m), 4.22 - 4.32 (1H, m), 3.19 - 3.29 (1H, m) 3.10 - 3.18 (1H, m) 2.46 (3H, s), 1.94 - 2.03 (1H, m); ^**13**^**C NMR** (101 MHz, DMSO-*d*_6_) δ = 168.2, 152.1, 146.9, 138.8, 131.5, 129.7, 128.9 (2C), 127.8 (2C), 68.4, 57.8, 52.3, 42.2, 38.2, 15.5; **LCMS** (Method A): t_R_ = 0.35 min, [M+H]^+^ 318, (95% purity); **HRMS** (C_16_H_20_N_3_O_2_S) [M+H]^+^ requires 318.1276, found [M+H]^+^ 318.1278.

#### *Tert*-butyl((*S*)-1-((2*S*,4*S*)-4-hydroxy-2-((4-(4-methylthiazol-5-yl)benzyl)carbamoyl)pyrrolidin-1-yl)-3,3-dimethyl-1-oxobutan-2-yl)carbamate (11)

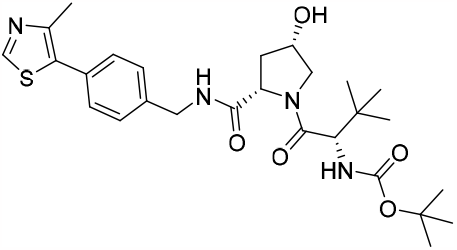

HATU (4.36 g, 11.48 mmol) was added to a stirred solution of (2*S*,4*S*)-4-hydroxy-*N*-(4-(4-methylthiazol-5-yl)benzyl)pyrrolidine-2-carboxamide, 2HCl (2.987 g, 7.65 mmol), DIPEA (4.01 mL, 22.96 mmol) and (*S*)-2- ((*tert*-butoxycarbonyl)amino)-3,3-dimethylbutanoic acid (1.770 g, 7.65 mmol) in DMF (15.31 mL) and the reaction mixture was stirred at room temperature for 1 h. The reaction mixture was diluted with water (75 mL) and extracted with DCM (3 × 75 mL). The organic layers were combined, passed through a hydrophobic frit and the solvent concentrated in vacuo. The residue was purified by reverse phase column chromatography (20 - 85% MeCN in H_2_O + 0.1% NH_4_HCO_3_, 330 g C18, 10 CV) to afford *tert*-butyl ((*S*)-1- ((2*S*,4*S*)-4-hydroxy-2-((4-(4-methylthiazol-5-yl)benzyl)carbamoyl)pyrrolidin-1-yl)-3,3-dimethyl-1-oxobutan-2-yl)carbamate (2.873 g, 5.41 mmol, 71 % yield) as an off-white foam. ^**1**^**H NMR** (400 MHz, DMSO-*d*_6_) δ = 8.99 (1H, s), 8.66 (1H, br t, *J* = 5.50 Hz), 7.37 - 7.43 (4H, m), 6.59 (1H, br d, *J* = 8.44 Hz), 5.44 (1H, br d, *J* = 6.97 Hz), 4.44 (1H, dd, *J* = 15.77, 6.60 Hz), 4.40 (1H, br dd, *J* = 8.44, 6.24 Hz) 4.27 (1H, dd, *J* = 15.77, 5.50 Hz), 4.21 - 4.24 (1H, m), 4.11 (1 H, br d, *J* = 8.44 Hz), 3.86 - 3.94 (1H, m), 3.41 - 3.47 (1H, m), 2.45 (3H, s), 2.32 - 2.38 (1H, m), 1.71 - 1.78 (1H, m), 1.38 (9H, s), 0.95 (9H, s); ^**13**^**C NMR** (151 MHz, DMSO-*d*_6_) δ = 172.4, 170.2, 155.6, 151.4, 147.7, 139.1, 131.1, 129.7, 128.7 (2C) 127.4 (2C), 78.1, 69.1, 58.5, 55.6, 54.9, 41.8, 36.8, 34.7, 28.2 (3C), 26.3 (3C), 15.9; **LCMS** (Method B): t_R_ = 1.04 min, [M+H]^+^ 532, (98% purity); **HRMS** (C_27_H_39_N_4_O_5_S) [M+H]^+^ requires 531.2641, found [M+H]^+^ 531.2642.

#### (3*S*,5*S*)-1-((*S*)-2-((*Tert*-butoxycarbonyl)amino)-3,3-dimethylbutanoyl)-5-((4-(4-methylthiazol-5-yl)benzyl)carbamoyl)-pyrrolidin-3-yl methanesulfonate (1)

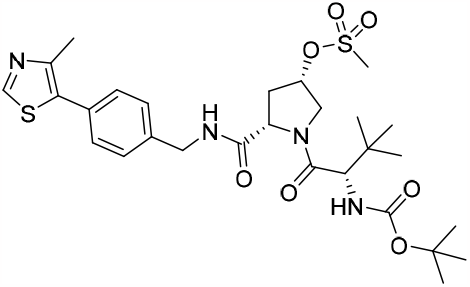

Mesyl chloride (0.501 mL, 6.44 mmol) was added dropwise to a stirred solution of *tert*-butyl ((*S*)-1-((2*S*,4*S*)-4-hydroxy-2-((4-(4-methylthiazol-5-yl)benzyl)carbamoyl)pyrrolidin-1-yl)-3,3-dimethyl-1-oxobutan-2-yl)carbamate (2.846 g, 5.36 mmol) and triethylamine (0.897 mL, 6.44 mmol) in DCM (17.88 mL) over an ice-water bath, and the reaction mixture was stirred at room temperature for 30 min under a nitrogen atmosphere. The reaction mixture was washed with 5% citric acid (50 mL) followed by water (50 mL) and the organic layer was passed through a hydrophobic frit, and the solvent removed in vacuo. The residue was purified by normal phase column chromatography (100% cyclohexane, 2 CV followed by 100% EtOAc, 330 g SiO_2_, 10 CV) to afford (3*S*,5*S*)-1-((*S*)-2-((*tert*-butoxycarbonyl)amino)-3,3-dimethylbutanoyl)-5-((4-(4-methylthiazol-5-yl)benzyl)carbamoyl)-pyrrolidin-3-yl methanesulfonate (2.954 g, 4.85 mmol, 90% yield) as a white foam. ^**1**^**H NMR** (400 MHz, DMSO-*d*_6_) δ = 8.99 (1H, s) 8.43 (1H, br t, *J* = 5.66 Hz) 7.37 - 7.44 (4H, m) 6.65 (1H, br d, *J* = 8.86 Hz) 5.23 - 5.36 (1H, m) 4.49 (1H, dd, *J* = 8.86, 5.91 Hz) 4.35 (2H, dd, *J* = 5.66, 2.71 Hz) 4.19 -4.28 (1H, m) 4.11 (1H, br d, *J* = 8.86 Hz) 3.71 (1H, dd, *J* = 11.32, 4.92 Hz) 3.23 (3H, s) 2.59 - 2.69 (1H, m) 2.45 (3H, s) 2.08 - 2.16 (1H, m) 1.39 (9H, s) 0.90 - 0.99 (9H, m); ^**13**^**C NMR** (101 MHz, DMSO-*d*_6_) δ = 170.4, 170.1, 155.6, 151.4, 147.7, 139.2, 131.1, 129.7, 128.7 (2C), 127.5 (2C), 78.1, 77.8, 59.7, 58.7, 57.8, 54.8, 52.6, 41.8, 37.6, 34.8, 28.1 (2C), 26.3 (2C), 15.9, 14.0; **LCMS** (Method A): t_R_ = 1.06 min, [M+H]^+^ 609, (100% purity); **HRMS** (C_28_H_40_N_4_O_7_S_2_) [M+H]^+^ requires 609.2417, found [M+H]^+^ 609.2415.

#### *S*-((3*R*,5*S*)-1-((*S*)-2-((*Tert*-butoxycarbonyl)amino)-3,3-dimethylbutanoyl)-5-((4-(4-methylthiazol-5-yl)benzyl)carbamoyl)pyrrolidin-3-yl) ethanethioate (2)

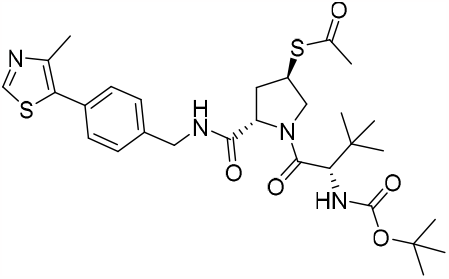

Thioacetic acid (0.445 mL, 6.20 mmol) was added to a stirred solution of Cs_2_CO_3_ (1.010 g, 3.10 mmol) and in DMF (11.55 mL) under a nitrogen atmosphere, and the reaction mixture was stirred at room temperature for 30 min within a sealed vessel. (3*S*,5*S*)-1-((*S*)-2-((*Tert*-butoxycarbonyl)amino)-3,3-dimethylbutanoyl)-5-((4-(4-methylthiazol-5-yl)benzyl)carbamoyl)pyrrolidin-3-yl methanesulfonate (2.904 g, 4.77 mmol) in DMF (4.35 mL) was added and the sealed reaction mixture was stirred at 50 °C for 24 h. The solvent was removed in vacuo and the residue was diluted with ethyl acetate (100 mL) washed with saturated aqueous sodium hydrogen carbonate (100 mL) followed by water (100 mL) and the organic layer was passed through a hydrophobic frit, and the solvent removed in vacuo. The residue was purified normal phase column chromatography (0 - 100% EtOAc in cyclohexane, 330 g SiO_2_, 10 CV) to afford *S*-((3*R*,5*S*)-1-((*S*)-2-((*tert*-butoxycarbonyl)amino)-3,3-dimethylbutanoyl)-5-((4-(4-methylthiazol-5-yl)benzyl)carbamoyl)pyrrolidin-3-yl) ethanethioate (2.435 g, 4.14 mmol, 87% yield) as an off-white solid. ^**1**^**H NMR** (400 MHz, DMSO-*d*_6_) δ = 8.99 (1H, s), 8.57 (1H, br t, *J* = 5.66 Hz), 7.38 - 7.41 (4H, m), 6.65 (1H, br d, *J* = 9.35 Hz), 4.47 (1H, br t, *J* = 7.14 Hz), 4.36 - 4.44 (1H, m), 4.22 - 4.32 (1H, m), 3.98 - 4.12 (3H, m), 3.71 - 3.80 (1H, m), 2.45 (3H, s), 2.34 (3H, s), 2.22 - 2.31 (1H, m), 2.10 - 2.19 (1H, m), 1.39 (9H, s), 0.94 (9H, s); ^**13**^**C NMR** (101 MHz, DMSO-*d*_6_) δ = 194.9, 171.0, 169.8, 155.6, 151.4, 147.7, 139.2, 131.1, 129.7, 128.7 (2C), 127.4 (2C), 78.1, 58.6, 58.5, 53.1, 41.6, 40.9, 34.7, 34.6, 30.5, 28.1 (3C), 26.2 (3C), 15.9; **LCMS** (Method A): t_R_ = 1.19 min, [M+H]^+^ 589, (98% purity); **HRMS** (C_29_H_41_N_4_O_5_S_2_) [M+H]^+^ requires 589.2518, found [M+H]^+^ 589.2515.

#### *Tert*-butyl ((2*S*)-1-((4*R*)-4-(fluorosulfonyl)-2-((4-(4-methylthiazol-5-yl)benzyl)carbamoyl)pyrrolidin-1-yl)- 3,3-dimethyl-1-oxobutan-2-yl)carbamate (4)

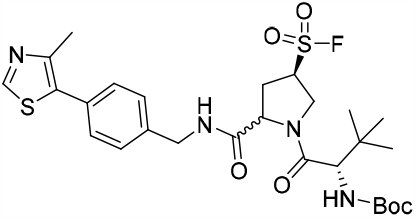

2 M aqueous HCl (42.5 μL, 0.085 mmol) was added dropwise to a stirred solution of NCS (91 mg, 0.679 mmol) in MeCN (233 uL) over an ice-water bath, and the reaction mixture was stirred over an ice-water bath for 15 min. *S*-((3*R*,5*S*)-1-((*S*)-2-((*Tert*-butoxycarbonyl)amino)-3,3-dimethylbutanoyl)-5-((4-(4-methylthiazol-5-yl)benzyl)carbamoyl)pyrrolidin-3-yl) ethanethioate (100 mg, 0.170 mmol) in MeCN (50 uL) was added dropwise and the reaction mixture was stirred at room temperature for 20 min. The reaction mixture was diluted with DCM (10 mL) washed with brine (3 × 5 mL) and the organic layer was passed through a hydrophobic frit and the solvent removed in vacuo to afford *tert*-butyl ((*S*)-1-((2*S*,4*R*)-4-(chlorosulfonyl)-2- ((4-(4-methylthiazol-5-yl)benzyl)carbamoyl)pyrrolidin-1-yl)-3,3-dimethyl-1-oxobutan-2-yl)carbamate (104 mg, 0.170 mmol, 100% yield) as a yellow oil. Potassium fluoride (39.4 mg, 0.678 mmol) and 18-crown-6 (179 mg, 0.678 mmol) were added to a stirred solution of *tert*-butyl ((*S*)-1-((2*S*,4*R*)-4-(chlorosulfonyl)-2-((4-(4-methylthiazol-5-yl)benzyl)carbamoyl)pyro-lidin-1-yl)-3,3-dimethyl-1-oxobutan-2-yl)carbamate (104 mg, 0.170 mmol) in MeCN (848 μL) under a nitrogen atmosphere, and the reaction mixture was sealed and stirred at room temperature 1 h. The reaction was abandoned due to racemisation of the proline ring.

#### *Tert*-butyl ((*S*)-1-((2*S*,4*R*)-4-(fluorosulfonyl)-2-((4-(4-methylthiazol-5-yl)benzyl)carbamoyl)pyrrolidin-1-yl)- 3,3-dimethyl-1-oxobutan-2-yl)carbamate (5)

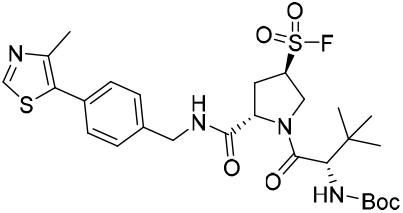

Selectfluor (451 mg, 1.274 mmol), *S*-((3*R*,5*S*)-1-((*S*)-2-((tert-butoxycarbonyl)amino)-3,3-dimethylbutanoyl)-5- ((4-(4-methylthiazol-5-yl)benzyl)carbamoyl)pyrrolidin-3-yl)ethanethioate (100 mg, 0.170 mmol) in acetonitrile (1544 μL) and water (154 μL) were sealed within a vessel and heated in a Biotage Initiator microwave for 15 min at 60 °C using a normal absorption setting. The reaction mixture was allowed to cool to room temperature. The reaction mixture was diluted with brine (2 mL) and extracted with DCM (3 × 10 mL). The organic layers were combined, passed through a hydrophobic frit and the solvent removed in vacuo. The residue was purified by normal phase column chromatography (0 - 100% EtOAc in cyclohexane, 40 g SiO_2_, 12 CV) to afford *tert*-butyl ((*S*)-1-((2*S*,4*R*)-4-(fluorosulfonyl)-2-((4-(4-methylthiazol-5-yl)benzyl)carbamoyl)pyrrolidin-1-yl)-3,3-dimethyl-1-oxobutan-2-yl)carbamate (38 mg, 0.064 mmol, 38% yield) as a white foam. ^**1**^**H NMR** (400 MHz, DMSO-*d*_6_) δ = 8.98 (1H, s), 8.69 (1H, br t, *J* = 5.91 Hz), 7.40 (4H, s), 6.77 (1H, br d, *J* = 9.35 Hz), 4.88 - 4.99 (1H, m), 4.62 (1H, br t, *J* = 7.88 Hz), 4.54 (1H, br d, *J* = 12.31 Hz), 4.42 (1H, dd, *J* = 15.75, 6.40 Hz), 4.23 - 4.30 (1H, m), 4.16 (2H, br d, *J* = 9.35 Hz), 2.65 - 2.77 (2H, m), 2.44 (3H, s), 1.38 (9H, s), 0.94 (9H, s); ^**13**^**C NMR** (151 MHz, DMSO-*d*_6_) δ = 170.3, 170.0, 155.8, 151.4, 147.7, 139.0, 131.1, 129.8, 128.7 (2C), 127.5 (2C), 78.3, 59.9 (br d, *J* = 14.37 Hz), 58.5, 58.3, 47.9, 41.7, 34.5, 30.1, 28.1, 26.2 (3C), 15.9; ^**19**^**F NMR** (376 MHz, DMSO-*d*_6_) δ = 47.66 (1F, s) **LCMS** (Method A): t_R_ = 1.18 min, [M+H]^+^ 497 (Boc-deprotected fragment), (99% purity); **HRMS** (C_27_H_38_FN_4_O_6_S_2_) [M+H]^+^ requires 597.2217, found [M+H]^+^ 597.2219.

#### (3*R*,5*S*)-1-((*S*)-2-Amino-3,3-dimethylbutanoyl)-5-((4-(4-methylthiazol-5-yl)benzyl)carbamoyl)-pyrrolidine-3-sulfonyl fluoride, 2HCl (6)

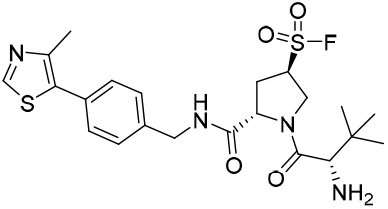

4 M HCl in 1,4-dioxane (356 μL, 1.424 mmol) was added to a stirred solution of *tert*-butyl ((*S*)-1-((2*S*,4*R*)-4- (fluorosulfonyl)-2-((4-(4-methylthiazol-5-yl)benzyl)carbamoyl)pyrrolidin-1-yl)-3,3-dimethyl-1-oxobutan-2-yl)carbamate (34 mg, 0.057 mmol) in 1,4-dioxane (114 μL) and the reaction mixture was stirred at room temperature 1 h. The solvent was removed in vacuo to afford (3*R*,5*S*)-1-((*S*)-2-amino-3,3-dimethylbutanoyl)- 5-((4-(4-methylthiazol-5-yl)benzyl)carbamoyl)pyrrolidine-3-sulfonyl fluoride, 2HCl (30 mg, 0.053 mmol, 92% yield) as a white solid. ^**1**^**H NMR** (400 MHz, DMSO-*d*_6_) δ = 9.01 (1H, s), 8.86 (1H, t, *J* = 5.91 Hz), 8.12 - 8.32 (4H, m), 7.41 (3H, d, *J* = 2.46 Hz), 4.99 - 5.08 (1H, m), 4.74 (1H, t, *J* = 7.88 Hz), 4.54 (1H, dd, *J* = 13.04, 2.71 Hz), 4.40 - 4.48 (1H, m), 4.28 (1H, dd, *J* = 15.75, 5.41 Hz), 4.08 - 4.13 (2H, m), 2.71 - 2.81 (1H, m), 2.46 (3H, s), 1.04 (9H, s); **LCMS** (Method A): t_R_ = 0.61 min, [M+H]^+^ 497, (98% purity); ^**19**^**F NMR** (376 MHz, DMSO-*d*_6_) δ = 48.59 (1F, s); **HRMS** (C_22_H_30_FN_4_O_4_S_2_) [M+H]^+^ requires 497.1693, found [M+H]^+^ 497.1695.

#### (3*R*,5*S*)-1-((*S*)-2-Acetamido-3,3-dimethylbutanoyl)-5-((4-(4-methylthiazol-5-yl)benzyl)carbamoyl)- pyrrolidine-3-sulfonyl fluoride (VHL-SF1)

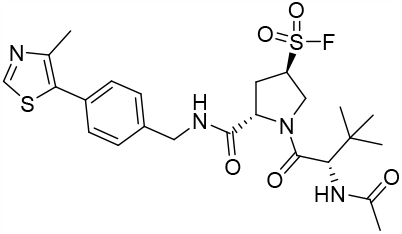

HATU (28 mg, 0.074 mmol) was added to a stirred solution of (3*R*,5*S*)-1-((*S*)-2-amino-3,3-dimethylbutanoyl)- 5-((4-(4-methylthiazol-5-yl)benzyl)carbamoyl)pyrrolidine-3-sulfonyl fluoride, 2HCl (28 mg, 0.049 mmol), DIPEA (25.8 μL, 0.147 mmol) and acetic acid (3.09 μL, 0.054 mmol) in DMF (98 μL) and the reaction mixture was stirred at room temperature for 30 min. The reaction mixture was diluted with brine (2 mL) and extracted with DCM (3 × 10 mL). The organic layers were combined, passed through a hydrophobic frit and the solvent removed in vacuo. The residue was purified directly by normal phase column chromatography (0 - 100% EtOAc in cyclohexane, 12 g SiO_2_, 10 CV) to afford (3*R*,5*S*)-1-((*S*)-2-acetamido-3,3-dimethylbutanoyl)-5-((4-(4- methylthiazol-5-yl)benzyl)carbamoyl)-pyrrolidine-3-sulfonyl fluoride (19 mg, 0.035 mmol, 72% yield) as a white solid. ^**1**^**H NMR** (400 MHz, DMSO-*d*_6_) δ = 8.98 (1H, s), 8.68 (1H, t, *J* = 5.91 Hz), 7.99 (1H, d, *J* = 8.86 Hz), 7.40 (4H, s), 4.90 - 4.99 (1H, m), 4.61 (1H, t, *J* = 7.63 Hz), 4.45 (2H, d, *J* = 8.86 Hz), 4.41 (1H, d, *J* = 6.40 Hz), 4.22 - 4.30 (1H, m), 4.13 - 4.21 (1H, m), 2.64 - 2.73 (2H, m), 2.44 (3H, s), 1.87 (3H, s), 0.96 (9H, s); ^**13**^**C NMR** (151 MHz, DMSO-*d*_6_) δ =170.3, 169.6, 169.4, 151.5, 147.7, 139.0, 131.1, 129.8, 128.7 (2C), 127.5 (2C), 59.7 (d, *J* = 14.38 Hz), 58.4, 56.7, 47.8, 41.7, 34.6, 30.1, 26.2 (3C), 22.0, 15.9; ^**19**^**F NMR** (376 MHz, DMSO-*d*_6_) δ = 47.61 (1F, s); **LCMS** (Method A): t_R_ = 0.90 min, [M+H]^+^ 539, (100% purity); **HRMS** (C_24_H_31_FN_4_O_5_S_2_) [M+H]^+^ requires 539.1798, found [M+H]^+^ 539.1794.

#### (3*R*,5*S*)-1-((*S*)-2-(*Tert*-butyl)-4,16-dioxo-20-((3a*S*,4*R*,6a*R*)-2-oxohexahydro-1*H*-thieno[3,4-d]imidazol-4-yl)- 6,9,12-trioxa-3,15-diazaicosanoyl)-5-((4-(4-methylthiazol-5-yl)benzyl)carbamoyl-)pyrrolidine-3-sulfonyl fluoride (VHL-SF1-Biotin)

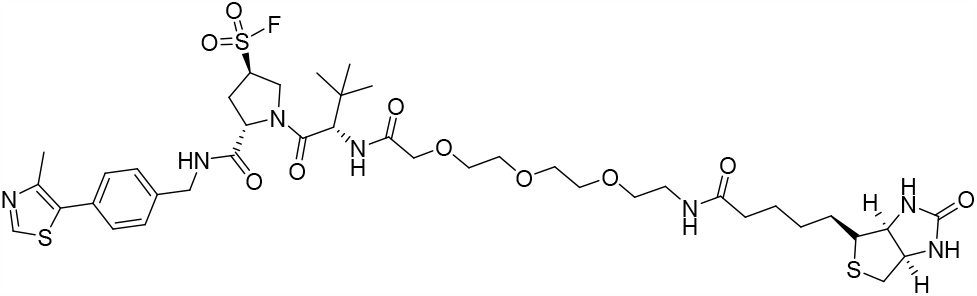

HATU (73.1 mg, 0.192 mmol) was added to a stirred solution of (3*R*,5*S*)-1-((*S*)-2-amino-3,3-dimethylbutanoyl)-5-((4-(4-methylthiazol-5-yl)benzyl)carbamoyl)pyrrolidine-3-sulfonyl fluoride, 2HCl (73 mg, 0.128 mmol), DIPEA (67.2 μL, 0.385 mmol) and 13-oxo-17-((3a*S*,4*R*,6a*R*)-2-oxohexahydro-1*H*-thieno[3,4-d]imidazol-4-yl)-3,6,9-trioxa-12-azaheptadecanoic acid (61.1 mg, 0.141 mmol) in DMF (256 μL) and the reaction mixture was stirred at room temperature for 30 min. The reaction mixture was diluted with water (2 mL) and extracted with DCM (3 × 10 mL). The organic layers were combined, passed through a hydrophobic frit and the solvent removed in vacuo. The residue was purified by normal phase column chromatography (0 - 100% EtOAc in cyclohexane, 24 g SiO_2_, 10 CV, followed by 0 -20% MeOH in EtOAc, 5 CV, 20% MeOH in EtOAc, 20 CV) to afford (3*R*,5*S*)-1-((*S*)-2-(*tert*-butyl)-4,16-dioxo-20-((3a*S*,4*R*,6a*R*)-2-oxohexahydro-1*H*- thieno[3,4-d]imidazol-4-yl)-6,9,12-trioxa-3,15-diazaicosanoyl)-5-((4-(4-methylthiazol-5-yl)benzyl)carbamoyl)pyrrolidine-3-sulfonyl fluoride (64 mg, 0.070 mmol, 55% yield) as a white foam. ^**1**^**H NMR** (400 MHz, DMSO-*d*_6_) δ = 8.99 (1H, s), 8.72 (1H, t, *J* = 5.91 Hz), 7.80 (1H, br t, *J* = 5.66 Hz), 7.45 - 7.49 (1H, m), 7.41 (1H, s), 7.38 - 7.45 (4H, m), 4.93 - 5.01 (1H, m), 4.64 (1H, t, *J* = 7.63 Hz), 4.56 (1H, d, *J* = 9.35 Hz), 4.42 - 4.47 (1H, m), 4.40 (1H, d, *J* = 6.40 Hz), 4.26 - 4.34 (2H, m), 4.16 - 4.24 (1H, m), 4.10 - 4.16 (2H, m), 3.97 (2H, s), 3.51 - 3.63 (9H, m), 3.37 - 3.41 (2H, m), 3.15 - 3.20 (2H, m), 3.06 - 3.12 (1H, m), 2.82 (2H, dd, *J* = 12.31, 5.41 Hz), 2.73 (1H, m), 2.46 (3H, s), 2.06 (2H, t, *J* = 7.38 Hz), 1.43 - 1.55 (4H, m), 1.25 - 1.34 (2H, m), 0.97 (9H, s); ^**13**^**C NMR** (151 MHz, DMSO-*d*_6_) δ = 172.1, 170.3, 170.1, 169.1, 169.0, 162.6, 151.5, 147.7, 139.0, 131.0, 129.8, 128.9, 128.7, 128.2, 127.5, 70.3, 69.7, 69.5, 69.5, 69.3, 69.1, 61.0, 59.7, 59.2, 58.3, 55.8, 55.4, 48.1, 41.8, 38.4, 35.0, 30.0, 28.2, 28.0, 26.0 (3C), 25.2, 15.9, 14.1; ^**19**^**F NMR** (376 MHz, DMSO-*d*_6_) δ = 47.79 (1F, s); **LCMS** (Method A): t_R_ = 0.88 min, [M+H]^+^ 912, (95% purity); **HRMS** (C_40_H_59_FN_7_O_10_S_3_) [M+H]^+^ requires 912.3470, found [M+H]^+^ 912.3456.

### Synthesis of VHL-SF2 and VHL-SF2-Biotin

#### *Tert*-butyl (*S*)-3-(4-bromophenyl)-3-((*tert*-butoxycarbonyl)-amino)propanoate (13)

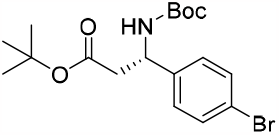

To a solution of (*S*)-3-(4-bromophenyl)-3-((*tert*-butoxycarbonyl)amino)propanoic acid (50 mg, 145.3 μmol) in toluene (484 μL) was added 1,1-ditert-butoxy-*N,N*-dimethyl-methanamine (118.1 mg, 139 μL, 581.1 μmol) and the sealed reaction mixture was heated at 130 °C for 40 min. The reaction mixture was allowed to cool to room temperature and diluted with DCM (25 mL), washed with saturated aqueous sodium bicarbonate (25 mL), and passed through a hydrophobic frit. The solvent was removed in vacuo to afford *tert*-butyl (*S*)-3- (4-bromophenyl)-3-((*tert*-butoxycarbonyl)-amino)propanoate (49 mg, 122.41 μmol, 84%) as a white solid. ^**1**^**H NMR** (400 MHz, DMSO-*d*_6_) δ = 7.51 (2H, d, *J* = 8.37 Hz), 7.44 - 7.50 (1H, m), 7.26 (2H, d, *J* = 8.37 Hz), 4.78 - 4.90 (1H, m), 2.55 - 2.63 (2H, m), 1.28 - 1.39 (18H, m); ^**13**^**C NMR** (151 MHz, DMSO-*d*_6_) δ = 169.3, 154.7, 142.2, 131.2 (2C), 128.8 (2C), 120.1, 80.1, 78.1, 50.9, 42.2, 28.2 (3C), 27.6 (3C); LCMS (Method A): t_R_ = 1.39 min, [M+H]^+^ 401 & 403, (98 % purity); **HRMS** (C_18_H_26_BrNO_4_Na) [M+Na]^+^ requires 422.0943, found [M+H]^+^ 422.0943.

#### *Tert*-butyl (*S*)-3-((tert-butoxycarbonyl)amino)-3-(4-(4-methylthiazol-5-yl)phenyl)propanoate (14)

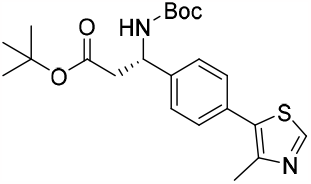

A stirred suspension of *tert*-butyl (*S*)-3-(4-bromophenyl)-3-((*tert*-butoxycarbonyl)amino)propanoate (5.815 g, 12.49 mmol), 4-methylthiazole (2.27 mL, 24.99 mmol), PdOAc_2_ (70 mg, 312.31 μmol), potassium carbonate (3.453 g, 24.99 mmol), pivalic acid (423 μL, 3.75 mmol) and DMA (41 mL) was heated at 130 °C for 16 h open to the atmosphere. The reaction mixture was filtered over Celite, diluted with saturated aqueous sodium hydrogen carbonate (100 mL), and extracted with DCM (3 × 100 mL). The organic layers were combined, passed through a hydrophobic frit and the solvent concentrated in vacuo. The residue was purified by normal phase column chromatography (0 - 50% ethyl acetate in cyclohexane, 330 g SiO_2_, 12 CV) to afford *tert*-butyl (*S*)-3-((*tert*-butoxycarbonyl)amino)-3-(4-(4-methylthiazol-5-yl)phenyl)propanoate (4.498 g, 10.75 mmol, 86%) as a yellow hard gum. ^**1**^**H NMR** (400 MHz, DMSO-*d*_6_) δ = 8.98 (1H, s), 7.52 (1H, br d, *J* = 9.35 Hz), 7.39 - 7.48 (4H, m), 4.88 - 4.98 (1H, m), 2.63 (2H, br d, *J* = 7.38 Hz), 2.45 (3H, s), 1.37 (9H, s), 1.34 (9H, s); **LCMS** (Method A): t_R_ = 1.27 min, [M+H]^+^ 401 & 403, (93 % purity).

#### *Tert*-butyl (*S*)-3-amino-3-(4-(4-methylthiazol-5-yl)phenyl)propanoate (15)

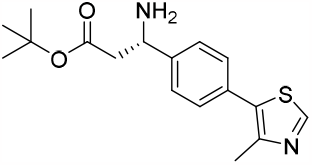

To a solution of *tert*-butyl (*S*)-3-((*tert*-butoxycarbonyl)amino)-3-(4-(4-methylthiazol-5-yl)phenyl)propanoate (1.34 g, 3.20 mmol) in MeOH (17 mL) was added 37% HCl (4.3 mL) dropwise and the reaction mixture was stirred at room temperature for 3 h. The reaction mixture was basified to pH 14 using conc. aqueous NaOH over an ice-water bath, and extracted with 10% MeOH in ethyl acetate (3 × 50 mL). The organic layers were combined, passed through a hydrophobic frit and the solvent removed in vacuo. The residue was purified by reverse phase column chromatography (5 - 65% MeCN in H_2_O + 0.1% NH_4_HCO_3_, 120 g C18, 12 CV) to afford *tert*-butyl (*S*)-3-amino-3-(4-(4-methylthiazol-5-yl)phenyl)propanoate (601 mg, 1.89 mmol, 59%) as a yellow oil. ^**1**^**H NMR** (400 MHz, DMSO-*d*_6_) δ = 8.98 (1H, s), 7.37 - 7.51 (4H, m), 4.19 (1H, t, *J* = 7.14 Hz), 2.53 - 2.56 (1H, m), 2.45 (3H, s), 2.03 (2H, br s), 1.33 (9H, s); **LCMS** (Method A): t_R_ = 1.03 min, [M+H]^+^ 319, (96% purity).

#### Methyl (2*S*,4*S*)-4-hydroxypyrrolidine-2-carboxylate (17)

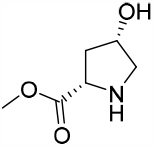

4 M HCl in Dioxane (40.8 mL, 163 mmol) was added to a solution of 1-(*tert*-butyl) 2-methyl (2*S*,4*S*)-4- hydroxypyrrolidine-1,2-dicarboxylate (10 g, 40.8 mmol) in DCM (163 mL) and MeOH (10 mL) the reaction mixture was stirred at room temperature 1 h. The solvent was removed in vacuo to afford methyl (2*S*,4*S*)-4-hydroxypyrrolidine-2-carboxylate, HCl (7.387 g, 40.7 mmol, 100% yield) as a white solid. ^**1**^**H NMR** (400 MHz, DMSO-*d*_6_) δ = 9.62 (1H, s), 5.41 (1H, d, *J* = 2.95 Hz), 4.50 (1H, dd, *J* = 9.84, 3.94 Hz), 4.33 - 4.42 (1H, m), 3.76 (3H, s), 3.19 - 3.25 (1H, m), 3.13 - 3.19 (1H, m), 2.27 - 2.39 (1H, m), 2.08 - 2.21 (1H, m); ^**13**^**C NMR** (101 MHz, DMSO-*d*_6_) δ = 169.6, 68.1, 57.3, 53.0, 52.9, 37.0.

#### Methyl (2S,4S)-4-hydroxy-1-(3-methyl-2-(3-methylisoxazol-5-yl)butanoyl)pyrrolidine-2-carboxylate (18)

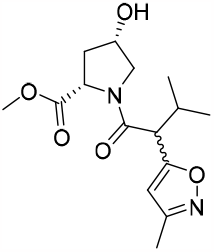

HATU (12.77 g, 33.6 mmol) was added to a stirred solution of methyl (2*S*,4*S*)-4-hydroxypyrrolidine-2-carboxylate, HCl (4.067 g, 22.39 mmol), DIPEA (11.73 mL, 67.2 mmol) and 3-methyl-2-(3-methylisoxazol-5-yl)butanoic acid (4.10 g, 22.39 mmol) in DCM (45 mL) and DMF (20mL) and the reaction mixture was stirred at room temperature for 1 h. The reaction mixture was diluted with saturated aqueous sodium hydrogen carbonate (75 mL) and extracted with DCM (3 × 75 mL). The organic layers were combined, passed through a hydrophobic frit and the solvent concentrated in vacuo. The residue was purified by reverse phase column chromatography (5 - 55% MeCN in H_2_O + 0.1% NH_4_HCO_3_, 130 g C18 (x3), 12 CV) to afford an orange oil. The residue was further purified by normal phase column chromatography (0 - 10% MeOH in TBME, 330 g SiO_2_, 10 CV) to afford methyl (2*S*,4*S*)-4-hydroxy-1-(3-methyl-2-(3-methylisoxazol-5-yl)butanoyl)pyrrolidine-2-carboxylate (3.867 g, 12.46 mmol, 56% yield) as light yellow oil. ^**1**^**H NMR** (400 MHz, DMSO-*d*_6_) δ = 6.16 - 6.26 (1H, m), 5.06 - 5.17 (1H, m), 4.93 - 5.03 (1H, m), 4.34 - 4.44 (1H, m), 4.25 - 4.33 (1H, m), 4.16 - 4.25 (1H, m), 3.60 - 3.68 (3H, m), 3.10 - 3.19 (1H, m), 2.25 - 2.37 (2H, m), 2.19 - 2.23 (3H, m), 1.78 - 1.86 (1H, m), 1.34 (3H, s), 1.12 (3H, s); **LCMS** (Method A): t_R_ = 0.66 min, [M+H]^+^ 311, (97% purity).

#### (2*S*,4*S*)-4-Hydroxy-1-((*R*)-3-methyl-2-(3-methylisoxazol-5-yl)butanoyl)pyrrolidine-2-carboxylic acid (19)

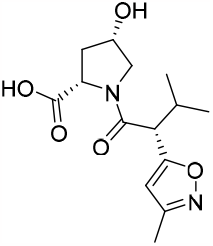

LiOH (0.597 g, 24.92 mmol) was added to a stirred solution of methyl (2*S*,4*S*)-4-hydroxy-1-(3-methyl-2-(3-methylisoxazol-5-yl)butanoyl)pyrrolidine-2-carboxylate (3.867 g, 12.46 mmol) in MeOH (8.31 mL) and water (4.15 mL) over an ice-water bath, and the reaction mixture was stirred for 2 h. The reaction mixture was diluted with water (40 mL) and extracted with EtOAc (3 × 40 mL) and the organic layers were discarded. The aqueous was acidified to pH 2 and extracted with 10:1 EtOAc:MeOH (4 × 50 mL). The organic layers were combined, passed through a hydrophobic frit and the solvent removed in vacuo. The enantiomers were separated (Chiralpak IE (250 × 30 mm, 5 μm), 70% Heptane (0.1% formic acid), 30% ethanol (0.1% formic acid); total flow rate: 40 mL/ min. Injection Volume: 400 μL; cycle time: 7.2 min on an Agilent 1200 Prep HPLCUV) to afford (2*S*,4*S*)-4-hydroxy-1-((*R*)-3-methyl-2-(3-methylisoxazol-5-yl)butanoyl)pyrrolidine-2-carboxylic acid (1.025 g, 29% yield) as a white solid. Absolute configuration determined by VCD. ^**1**^**H NMR** (400 MHz, DMSO-*d*_6_) δ = 6.08 - 6.26 (1H, m), 4.21 - 4.26 (1H, m), 3.76 - 3.81 (1H, m), 3.68 - 3.76 (1H, m), 3.44 (1H, dd, *J* = 10.27, 4.89 Hz), 3.22 - 3.29 (1H, m), 2.24 - 2.34 (2H, m), 2.17 - 2.20 (3H, m), 1.77 - 1.84 (1H, m), 0.87 - 1.04 (3H, m), 0.74 - 0.82 (3H, m); ^**13**^**C NMR** (101 MHz, DMSO-*d*_6_) δ = 172.6, 169.7, 167.7, 159.4, 103.2, 68.5, 57.3, 54.6, 49.0, 37.0, 31.1, 20.4, 19.8, 10.9; **LCMS** (Method A): t_R_ = 0.59 min, [M+H]^+^ 297, (98% purity); **HRMS** (C_14_H_21_N_2_O_5_) [M+H]^+^ requires 297.1450, found [M+H]^+^ 297.1455.

#### *Tert*-butyl (*S*)-3-((2*S*,4*S*)-4-hydroxy-1-((*R*)-3-methyl-2-(3-methylisoxazol-5-yl)butanoyl)pyrrolidine-2-carboxamido)-3-(4-(4-methylthiazol-5-yl)phenyl)propanoate (20)

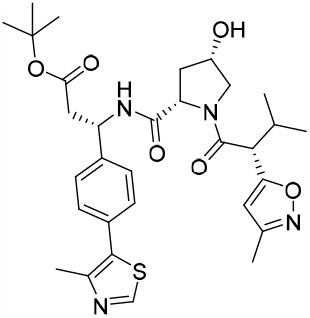

HATU (1.07 g, 2.82 mmol) was added to a stirred solution of (2*S*,4*S*)-4-hydroxy-1-((*R*)-3-methyl-2-(3-methylisoxazol-5-yl)butanoyl)pyrrolidine-2-carboxylic acid (556 mg, 1.88 mmol), DIPEA (654 μL, 3.75 mmol) and *tert*-butyl (*S*)-3-amino-3-(4-(4-methylthiazol-5-yl)phenyl)propanoate (598 mg, 1.88 mmol) in DMF (4 mL) and the reaction mixture was stirred at room temperature for 1 h. The reaction mixture was diluted with water (30 mL) and extracted with DCM (3 × 30 mL). The organic layers were combined, passed through a hydrophobic frit and the solvent removed in vacuo. The residue was purified by reverse phase column chromatography (30 - 95% MeCN in H_2_O + 0.1% HCOOH, 100 g C18, 12 CV). The volatile solvents were removed in vacuo and the aqueous neutralised to pH 7 using 2 M NaOH. The aqueous was extracted with DCM (3 × 100 mL), the organic layers were combined, passed through a hydrophobic frit and the solvent removed in vacuo to afford *tert*-butyl (*S*)-3-((2*S*,4*S*)-4-hydroxy-1-((*R*)-3-methyl-2-(3-methylisoxazol-5-yl)butanoyl)pyrrolidine-2-carboxamido)-3-(4-(4-methylthiazol-5-yl)phenyl)propanoate (846 mg, 1.42 mmol, 76%) as a hard yellow gum. ^**1**^**H NMR** (400 MHz, DMSO-*d*_6_) δ = 8.99 (1H, s), 8.43 (1H, d, *J* = 8.31 Hz), 7.38 - 7.47 (4H, m), 6.22 (1H, s), 5.26 (1H, br s), 4.26 (1H, dd, *J* = 8.93, 5.26 Hz), 4.16 (1H, br d, *J* = 3.42 Hz), 3.81 (1H, d, *J* = 9.54 Hz), 3.68 (1H, dd, *J* = 10.27, 5.38 Hz), 3.51 (1H, dd, *J* = 10.15, 4.52 Hz), 3.27 (1H, s), 2.74 - 2.83 (1H, m), 2.63 - 2.72 (1H, m), 2.45 (3H, s), 2.22 - 2.32 (2H, m), 2.19 (3H, s), 1.61 (1H, m), 1.31 (9H, s), 0.95 (3H, d, *J* = 6.60 Hz), 0.77 (3H, d, *J* = 6.60 Hz); **LCMS** (Method A): t_R_ = 1.09 min, [M+H]^+^ 597, (100% purity).

#### *Tert*-butyl (*S*)-3-((2*S*,4*S*)-1-((*R*)-3-methyl-2-(3-methylisoxazol-5-yl)butanoyl)-4-((methylsulfonyl)- oxy)pyrrolidine-2-carboxamido)-3-(4-(4-methylthiazol-5-yl)phenyl)propanoate (21)

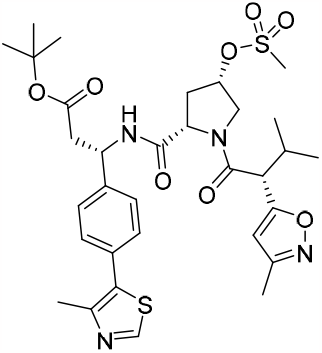

Mesyl chloride (351 μL, 4.50 mmol) was added dropwise to a stirred solution of *tert*-butyl (*S*)-3-((2*S*,4*S*)-4-hydroxy-1-((*R*)-3-methyl-2-(3-methylisoxazol-5-yl)butanoyl)pyrrolidine-2-carboxamido)-3-(4-(4-methylthiazol-5-yl)phenyl)propanoate (2.24 g, 3.75 mmol) and triethylamine (627 μL, 4.50 mmol) in DCM (13 mL) over an ice-water bath, and the reaction mixture was stirred at room temperature for 30 min under a nitrogen atmosphere. The reaction mixture was washed with 5 % citric acid (50 mL) followed by water (50 mL) and the organic layer was passed through a hydrophobic frit and the solvent removed in vacuo. The residue was purified by normal phase column chromatography (100% cyclohexane, 2 CV followed by 100% EtOAc, 120 g SiO_2_, 10 CV) to afford *tert*-butyl (*S*)-3-((2*S*,4*S*)-1-((*R*)-3-methyl-2-(3-methylisoxazol-5-yl)butanoyl)-4-((methylsulfonyl)oxy)pyro-lidine-2-carboxamido)-3-(4-(4-methylthiazol-5-yl)phenyl)propanoate (2.237 g, 3.32 mmol, 88%) as a viscous yellow oil. **LCMS** (Method A): t_R_ = 1.12 min, [M-H]^+^ 673, (99% purity).

#### *Tert*-butyl (*S*)-3-((2*S*,4*R*)-4-(acetylthio)-1-((*R*)-3-methyl-2-(3-methylisoxazol-5-yl)butanoyl)pyrro-lidine--2-carboxamido)-3-(4-(4-methylthiazol-5-yl)phenyl)propanoate (22)

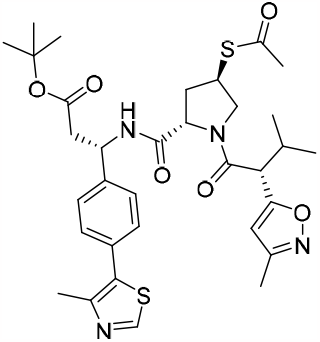

Thioacetic acid (119.8 μL, 1.67 mmol) was added to a stirred solution of Cs_2_CO_3_ (272 mg, 834.14 μmol) and in DMF (8.5 mL) under a nitrogen atmosphere, and the reaction mixture was stirred at room temperature for 30 min within a sealed vessel. *Tert*-butyl (*S*)-3-((2*S*,4*S*)-1-((*R*)-3-methyl-2-(3-methylisoxazol-5-yl)butanoyl)-4- ((methylsulfonyl)oxy)pyrrolidine-2-carboxamido)-3-(4-(4-methylthiazol-5-yl)phenyl)propanoate (866 mg, 1.28 mmol) in DMF (4 mL) was added and the sealed reaction mixture was stirred at 50 °C for 16 h. The solvent was removed in vacuo and the residue was purified by reverse phase column chromatography (40 - 95% MeCN in H_2_O + 0.1% NH_4_HCO_3_, 130 g C18, 15 CV) to afford *tert*-butyl (*S*)-3-((2*S*,4*R*)-4-(acetylthio)-1-((*R*)- 3-methyl-2-(3-methylisoxazol-5-yl)butanoyl)pyrrolidine-2-carboxamido)-3-(4-(4-methylthiazol-5-yl)phenyl)propanoate (368 mg, 561.97 mmol, 44%) as a yellow solid. ^**1**^**H NMR** (400 MHz, DMSO-*d*_6_) δ = 8.99 (1H, s), 8.50 (1H, d, *J* = 8.31 Hz), 7.37 - 7.47 (4H, m), 6.20 (1H, s), 5.09 - 5.20 (1H, m), 4.36 (1H, dd, *J* = 7.95, 5.75 Hz), 4.10 (1H, dd, *J* = 10.64, 6.48 Hz), 3.96 (1H, m), 3.79 (1H, d, *J* = 9.78 Hz), 3.46 (1H, dd, *J* = 10.64, 5.50 Hz), 2.64 - 2.83 (3H, m), 2.45 (3H, s), 2.30 (3H, s), 2.20 (3H, s), 2.01 - 2.15 (2H, m), 1.33 (9H, s), 0.97 (3H, d, *J* = 6.60 Hz), 0.77 (3H, d, *J* = 6.85 Hz); **LCMS** (Method A): t_R_ = 1.22 min, [M-H]^+^ 673, (97% purity).

#### *Tert*-butyl (*S*)-3-((2*S*,4R)-4-(fluorosulfonyl)-1-((*R*)-3-methyl-2-(3-methylisoxazol-5-yl)butanoyl)pyrr-olidine-2-carboxamido)-3-(4-(4-methylthiazol-5-yl)phenyl)propanoate (VHL-SF2)

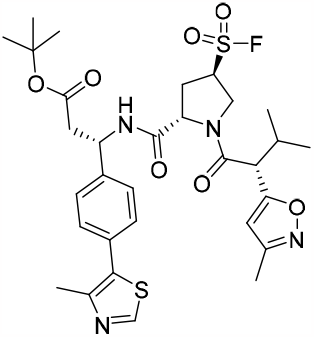

*Tert*-butyl (*S*)-3-((2*S*,4*R*)-4-(acetylthio)-1-((*R*)-3-methyl-2-(3-methylisoxazol-5-yl)butanoyl)pyrrolidine-2-carboxamido)-3-(4-(4-methylthiazol-5-yl)phenyl)propanoate (793 mg, 908.24 μmol) and selectfluor (2.413 g, 6.81 mmol) was added acetonitrile (8.256 mL) and water (825.67 μL) and the reaction mixture was sealed within a vessel and heated within a Biotage microwave for 20 min at 60 °C using a high absorption setting. The reaction mixture was cooled to room temperature. The reaction mixture was diluted with brine (10 mL) and extracted with DCM (3 × 10 mL), passed through a hydrophobic frit, and the solvent was removed in vacuo. The residue was purified by reverse phase column chromatography (40 - 95% MeCN + 0.1% HCO_2_H in H_2_O + 0.1% HCO_2_H, 100 g C18, 15 CV). The volatiles was removed in vacuo and the aqueous was neutralized with saturated aqueous sodium hydrogen carbonate and extracted with DCM (3 × 100 mL). The organic layers were combined, passed through a hydrophobic frit and the solvent removed in vacuo to afford *tert*-butyl (*S*)- 3-((2*S*,4R)-4-(fluorosulfonyl)-1-((*R*)-3-methyl-2-(3-methylisoxazol-5-yl)butanoyl)pyrrolidine-2-carboxamido)- 3-(4-(4-methylthiazol-5-yl)phenyl)propanoate (268 mg, 404.35 μmol, 45%) as a yellow foam. ^**1**^**H NMR** (400 MHz, DMSO-*d*_6_) δ = 8.99 (1H, s), 8.66 (1H, d, *J* = 8.07 Hz), 7.39 - 7.53 (4H, m), 6.21 (1H, s), 5.75 (1H, s), 5.08 - 5.19 (1H, m), 4.81 - 4.94 (1H, m), 4.50 - 4.58 (1H, m), 4.15 - 4.23 (1H, m), 3.96 (1H, d, *J* = 9.54 Hz), 2.66 - 2.85 (3H, m), 2.46 (3H, s), 2.22 - 2.34 (2H, m), 2.20 (3H, s), 1.34 (9H, s), 0.91 - 1.00 (3H, m), 0.80 (3H, d, *J* = 6.60 Hz); ^**13**^**C NMR** (101 MHz, DMSO-*d*_6_) δ = 169.7, 169.6, 169.6, 169.5, 168.0, 159.8, 152.0, 148.4, 141.9, 131.4, 130.8, 129.3 (2C), 127.6 (2C), 103.8, 80.8, 55.3, 50.2, 49.2, 48.2, 42.2, 31.4, 30.5, 28.1 (3C), 21.0, 20.2, 16.4, 11.4; ^**19**^**F NMR** (376 MHz, DMSO-*d*_6_) δ = 47.75 (1F, s); **LCMS** (Method A): t_R_ = 1.24 min, [M+H]^+^ 663, (99% purity); **HRMS** (C_31_H_40_FN_4_O_7_S_2_) [M+H]^+^ requires 663.2322, found [M+H]^+^ 663.2311.

#### (*S*)-3-((2*S*,4*R*)-4-(Fluorosulfonyl)-1-((*R*)-3-methyl-2-(3-methylisoxazol-5-yl)butanoyl)pyrrolidine-2- carboxamido)-3-(4-(4-methylthiazol-5-yl)phenyl)propanoic acid, 2TFA (23)

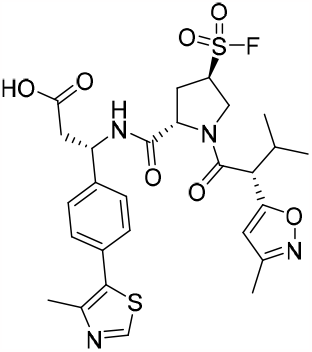

*Tert*-butyl (*S*)-3-((2*S*,4*R*)-4-(fluorosulfonyl)-1-((*R*)-3-methyl-2-(3-methylisoxazol-5-yl)butanoyl)pyrro-lidine-2-carboxamido)-3-(4-(4-methylthiazol-5-yl)phenyl)propanoate (50 mg, 75.44 μmol) in DCM (150.88 μL) was added TFA (75 μL, 973.5 μmol) and the reaction mixture was stirred at room temperature for 30 min. The solvent was removed in vacuo to afford (*S*)-3-((2*S*,4*R*)-4-(fluorosulfonyl)-1-((*R*)-3-methyl-2-(3-methylisoxazol-5-yl)butanoyl)pyrrolidine-2-carboxamido)-3-(4-(4-methylthiazol-5-yl)phenyl)propanoic acid, 2TFA (62 mg, 74.28 μmol, 99%) as an orange gum. ^**1**^**H NMR** (400 MHz, DMSO-*d*_6_) δ = 8.97 - 9.04 (1H, m), 7.86 - 7.94 (1H, m), 7.37 - 7.54 (4H, m), 6.22 (1H, s), 5.12 - 5.21 (1H, m), 4.47 - 4.57 (1H, m), 4.17 - 4.23 (1H, m), 3.96 (1H, d, *J* = 9.54 Hz), 2.77 - 2.86 (3H, m), 2.66 - 2.75 (3H, m), 2.32 - 2.35 (1H, m), 2.25 - 2.31 (1H, m), 2.20 (3H, s), 1.72 - 1.83 (3H, m), 0.92 - 1.03 (3H, m), 0.76 - 0.84 (3H, m); ^**19**^**F NMR** (376 MHz, DMSO-*d*_6_) δ = 47.80 (1F, s); **LCMS** (Method A): t_R_ = 0.77 min, [M+H]^+^ 607, (89% purity).

#### (3*R*,5*S*)-1-((*R*)-3-Methyl-2-(3-methylisoxazol-5-yl)butanoyl)-5-(((*S*)-1-(4-(4-methylthiazol-5-yl)phenyl)- 3,17-dioxo-21-((3a*S*,4*R*,6a*R*)-2-oxohexahydro-1*H*-thieno[3,4-d]imidazol-4-yl)-7,10,13-trioxa-4,16- diazahenicosyl)carbamoyl)pyrrolidine-3-sulfonyl fluoride (VHL-SF2-Biotin)

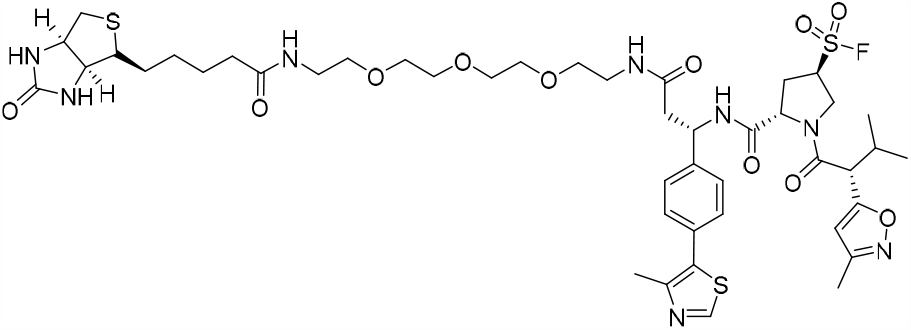

To a stirred solution of (*S*)-3-((2*S*,4*R*)-4-(fluorosulfonyl)-1-((*R*)-3-methyl-2-(3-methylisoxazol-5-yl)butanoyl)pyrrolidine-2-carboxamido)-3-(4-(4-methylthiazol-5-yl)phenyl)propanoic acid, 2TFA (60 mg, 35.94 μmol) and *N*-(2-(2-(2-(2-aminoethoxy)ethoxy)ethoxy)ethyl)-5-((3a*S*,4*R*,6a*R*)-2-oxohexahydro-1*H*- thieno[3,4-d]imidazol-4-yl)pentanamide, HCl (18 mg, 39.53 μmol) in DMF (72 μL) and DIPEA (18.8 μL, 107.82 μmol) was added HATU (21 mg, 53.91 μmol) and the reaction mixture was stirred at room temperature for 30 min. The reaction mixture was purified directly by MDAP (formic) to afford (3*R*,5*S*)-1-((*R*)-3-methyl-2-(3-methylisoxazol-5-yl)butanoyl)-5-(((*S*)-1-(4-(4-methylthiazol-5-yl)phenyl)-3,17-dioxo-21-((3a*S*,4*R*,6a*R*)-2-oxohexahydro-1*H*-thieno[3,4-d]imidazol-4-yl)-7,10,13-trioxa-4,16-diazahenicosyl)carbamoyl)pyrrolidine-3-sulfonyl fluoride (9 mg, 8.94 μmol, 25%) as a yellow solid. ^**1**^**H NMR** (400 MHz, DMSO-*d*_6_) δ = 8.98 (1H, s), 8.63 (1H, br d, *J* = 8.07 Hz), 7.97 (1H, br t, *J* = 5.87 Hz), 7.79 (1H, br t, *J* = 5.75 Hz), 7.40 - 7.48 (2H, m), 7.34 - 7.40 (2H, m), 6.31 - 6.42 (2H, m), 6.21 (1H, s), 5.15 - 5.24 (1H, m), 4.82 - 4.95 (2H, m), 4.48 - 4.56 (1H, m), 4.26 - 4.34 (1H, m), 4.20 (1H, br d, *J* = 4.89 Hz), 4.09 - 4.15 (1H, m), 3.96 (1H, br d, *J* = 9.29 Hz), 3.41 - 3.51 (6H, m), 3.38 (2H, t, *J* = 5.99 Hz), 3.12 - 3.22 (3H, m), 3.09 (1H, m), 2.81 (1H, dd, *J* = 12.35, 5.26 Hz), 2.65 - 2.75 (2H, m), 2.54 - 2.64 (3H, m), 2.42 - 2.48 (3H, m), 2.23 - 2.34 (2H, m), 2.15 - 2.22 (3H, m), 2.06 (2H, t, *J* = 7.34 Hz), 1.55 - 1.67 (2H, m), 1.40 - 1.54 (3H, m), 1.22 - 1.37 (5 H, m), 0.97 (3H, d, *J* = 6.60 Hz), 0.78 - 0.83 (3H, m); ^**19**^**F NMR** (376 MHz, DMSO-*d*_6_) δ = 47.78 (1F, s); **LCMS** (Method A): t_R_ = 0.90 min, [M+H]^+^ 589, (93% purity); **HRMS** (C_45_H_64_FN_8_O_11_S_3_) [M+H]^+^ requires 1007.3841, found [M+H]^+^ 1007.3846.

### Synthesis of BRD-SF2

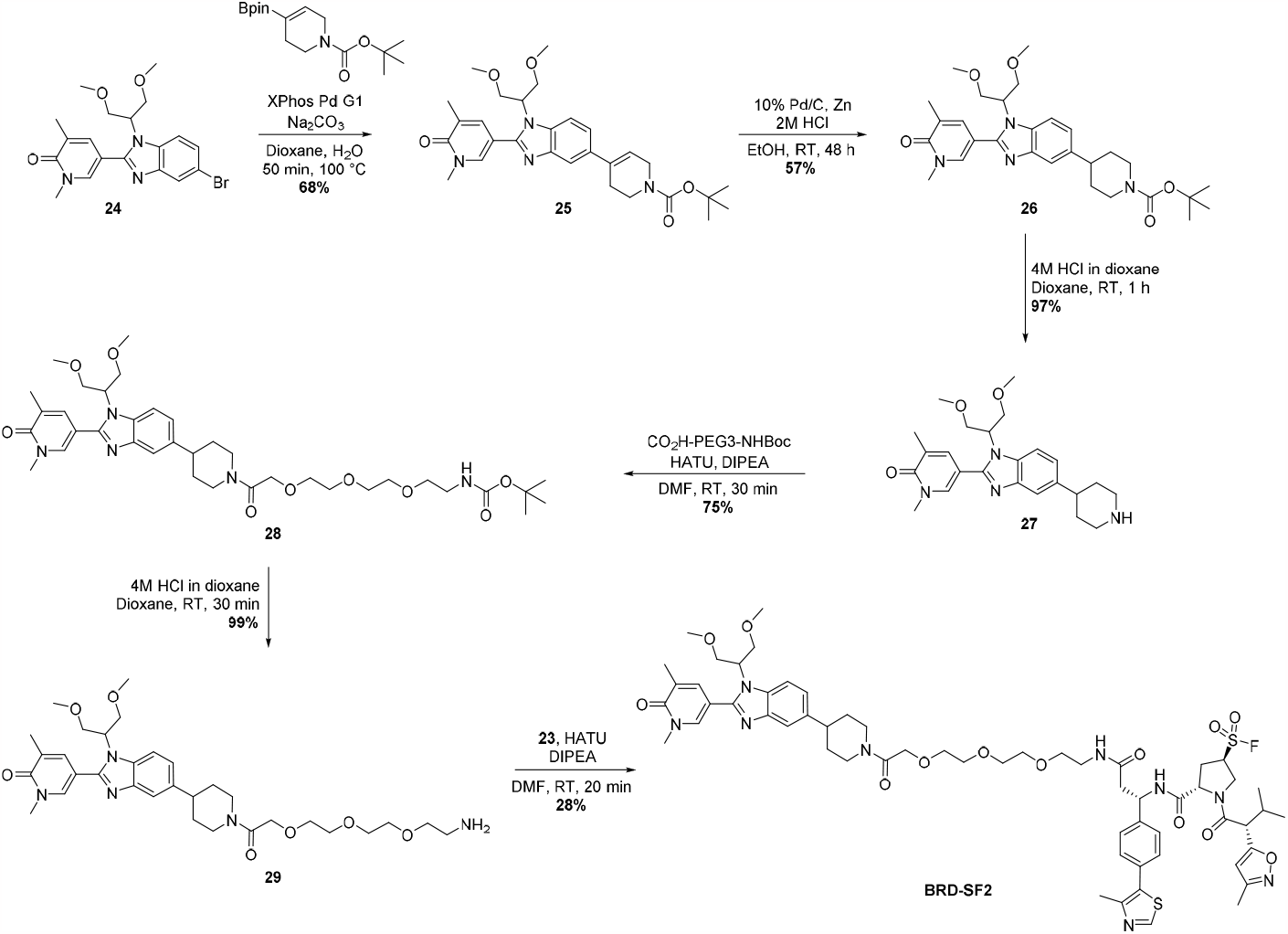

Benzimidazole **24** was synthesized according to literature precedent.^1^

#### *Tert*-butyl 4-(1-(1,3-dimethoxypropan-2-yl)-2-(1,5-dimethyl-6-oxo-1,6-dihydropyridin-3-yl)-1*H*-benzo[d]imidazol-5-yl)-3,6-dihydropyridine-1(2*H*)-carboxylate (25)

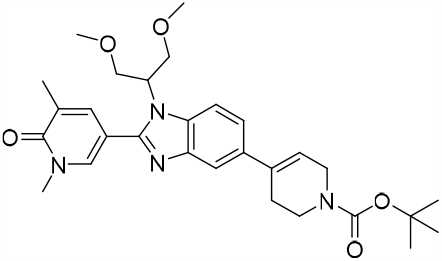

5-(5-Bromo-1-(1,3-dimethoxypropan-2-yl)-1*H*-benzo[d]imidazol-2-yl)-1,3-dimethylpyridin-2(1*H*)-one (200 mg, 475.84 μmol), *tert*-butyl 4-(4,4,5,5-tetramethyl-1,3,2-dioxaborolan-2-yl)-3,6-dihydropyridine-1(2*H*)- carboxylate (176.6 mg, 571.01 μmol), chloro(2-dicyclohexylphosphino-2’,4’,6’-triisopropyl-1,1’-biphenyl)[2- (2-aminoethyl)phenyl]palladium(II) (28.1 mg, 38.07 μmol), sodium carbonate (151.3 mg, 1.43 mmol) in water (332.8 μL) and 1,4-dioxane (1.331 mL) were sealed within a vessel, and the vessel was evacuated and purged three times with nitrogen. The reaction mixture was heated in a Biotage microwave for 50 min at 100 °C using a normal absorption setting. The reaction mixture was purified directly by MDAP (HpH) to afford *tert*-butyl 4-(1-(1,3-dimethoxypropan-2-yl)-2-(1,5-dimethyl-6-oxo-1,6-dihydropyridin-3-yl)-1*H*-benzo[d]imidazol-5-yl)-3,6-dihydropyridine-1(2*H*)-carboxylate (169 mg, 323.35 μmol, 68%) as a white solid. ^**1**^**H NMR** (400 MHz, DMSO-*d*_6_) δ = 8.99 (1H, s), 8.43 (1H, d, *J* = 8.31 Hz), 7.38 - 7.49 (4H, m), 5.26 (1H, br s), 5.16 (1H, q, *J* = 7.74 Hz), 4.26 (1H, dd, *J* = 8.93, 5.26 Hz), 4.13 - 4.22 (1H, m), 3.81 (1H, d, *J* = 9.54 Hz), 3.68 (1H, dd, *J* = 10.27, 5.38 Hz), 3.51 (1H, dd, *J* = 10.15, 4.52 Hz), 3.27 (1H, s), 2.75 - 2.83 (1H, m), 2.64 - 2.72 (1H, m), 2.45 (3H, s), 2.22 - 2.32 (2H, m), 2.19 (3H, s), 1.61 (1H, dt, *J* = 12.84, 5.32 Hz), 1.31 (9H, s); **LCMS** (Method B): t_R_ = 1.55 min, [M+H]^+^ 523, (99% purity); **HRMS** (C_29_H_39_N_4_O_5_) [M+H]^+^ requires 523.2920, found [M+H]^+^ 523.2919.

#### *Tert*-butyl 4-(1-(1,3-dimethoxypropan-2-yl)-2-(1,5-dimethyl-6-oxo-1,6-dihydropyridin-3-yl)-1*H*- benzo[d]imidazol-5-yl)piperidine-1-carboxylate (26)

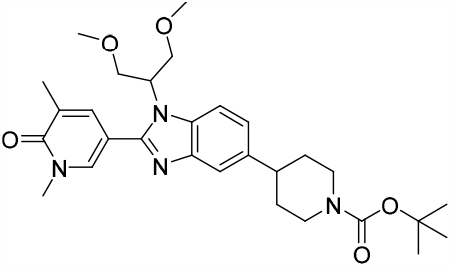

To a COware tube, 10% Pd/C (34 mg, 31.95 μmol) and *tert*-butyl 4-(1-(1,3-dimethoxypropan-2-yl)-2-(1,5-dimethyl-6-oxo-1,6-dihydropyridin-3-yl)-1*H*-benzo[d]imidazol-5-yl)-3,6-dihydropyridine-1(2*H*)-carboxylate (167 mg, 319.53 μmol) in ethanol (3.20 mL) was added to one tube under nitrogen and that tube was sealed. 2 M Aqueous HCl (639 μL, 1.28 mmol) and zinc (209 mg, 3.20 mmol) was added under nitrogen to the other tube which was subsequently sealed. The reaction mixture stirred at room temperature for 48 h. The reaction mixture was filtered over celite and washed with DCM (20 mL). The solvent was removed in vacuo and the sample was purified by MDAP (HpH) to afford *tert*-butyl 4-(1-(1,3-dimethoxypropan-2-yl)-2-(1,5-dimethyl-6-oxo-1,6-dihydropyridin-3-yl)-1*H*-benzo[d]imidazol-5-yl)piperidine-1-carboxylate (95 mg, 181.07 μmol, 57%) as a white solid. ^**1**^**H NMR** (400 MHz, DMSO-*d*_6_) δ = 8.01 (1H, d, *J* = 2.20 Hz), 7.69 (1H, d, *J* = 8.31 Hz), 7.65 (1H, dd, *J* = 2.45, 1.22 Hz), 7.46 (2H, d, *J* = 1.47 Hz), 7.11 (1H, dd, *J* = 8.56, 1.47 Hz), 4.81 (1H, tt, *J* = 8.80, 4.40 Hz), 4.10 (2H, br d, *J* = 12.23 Hz), 3.96 - 4.05 (2H, m), 3.75 (2H, dd, *J* = 10.51, 4.65 Hz), 3.54 (3H, s), 3.16 (6H, s), 2.76 - 2.85 (2H, m), 2.09 (3H, s), 1.80 (1H, s), 1.81 (2H, br d, *J* = 12.47 Hz), 1.49 - 1.62 (2H, m), 1.43 (9H, s); **LCMS** (Method B): t_R_ = 1.16 min, [M+H]^+^ 525, (100% purity); **HRMS** (C_29_H_41_N_4_O_5_) [M+H]^+^ requires 525.3077, found [M+H]^+^ 525.3080.

#### 5-(1-(1,3-Dimethoxypropan-2-yl)-5-(piperidin-4-yl)-1*H*-benzo[d]imidazol-2-yl)-1,3-dimethylpyridin-2(1*H*)- one, 2HCl (27)

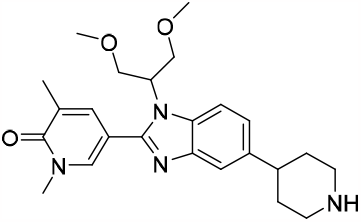

To a stirred solution of *tert*-butyl 4-(1-(1,3-dimethoxypropan-2-yl)-2-(1,5-dimethyl-6-oxo-1,6-dihydropyridin- 3-yl)-1*H*-benzo[d]imidazol-5-yl)piperidine-1-carboxylate (93 mg, 177.26 μmol) in 1,4-dioxane (354.5 μL) was added 4 M HCl in dioxane (1.11 mL, 4.43 mmol) and the reaction mixture was stirred at room temperature for 1 h. The solvent was removed by in vacuo to afford 5-(1-(1,3-dimethoxypropan-2-yl)-5-(piperidin-4-yl)- 1*H*-benzo[d]imidazol-2-yl)-1,3-dimethylpyridin-2(1*H*)-one, 2HCl (88 mg, 172.05 μmol, 97%) as a beige solid. ^**1**^**H NMR** (400 MHz, DMSO-*d*_6_) δ = 8.84 - 9.09 (2H, m), 8.26 (1H, br s), 8.06 (1H, br d, *J* = 8.31 Hz), 7.71 (1H, dd, *J* = 2.45, 0.98 Hz), 7.60 (1H, s), 7.36 (1H, br d, *J* = 8.56 Hz), 4.97 - 5.11 (1H, m), 4.09 (3H, br dd, *J* = 10.51, 9.05 Hz), 3.82 (3H, br dd, *J* = 10.64, 4.28 Hz), 3.36 - 3.46 (2H, m), 3.13 - 3.27 (6H, m), 2.95 - 3.12 (3H, m), 2.12 (3H, s), 1.95 - 2.05 (3H, m); **LCMS** (Method B): t_R_ = 0.76 min, [M+H]^+^ 425, (100% purity); **HRMS** (C_24_H_32_N_4_O_3_) [M+H]^+^requires 425.2553, found [M+H]^+^ 425.2557.

#### *Tert*-butyl (2-(2-(2-(2-(4-(1-(1,3-dimethoxypropan-2-yl)-2-(1,5-dimethyl-6-oxo-1,6-dihydropyridin-3-yl)- 1*H*-benzo[d]imidazol-5-yl)piperidin-1-yl)-2-oxoethoxy)ethoxy)ethoxy)ethyl)carbamate (28)

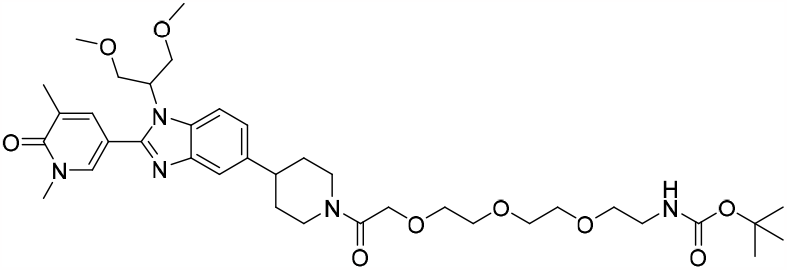

To a stirred solution of 5-(1-(1,3-dimethoxypropan-2-yl)-5-(piperidin-4-yl)-1*H*-benzo[d]imidazol-2-yl)-1,3- dimethylpyridin-2(1*H*)-one, 2HCl (86 mg, 172.87 μmol) and 2,2-dimethyl-4-oxo-3,8,11,14-tetraoxa-5-azahexadecan-16-oic acid (64 mg, 207.45 μmol) in DMF (346 μL) and DIPEA (60.2 μL, 345.75 μmol) was added HATU (132 mg, 345.75 μmol) and the reaction mixture was stirred at room temperature for 30 min. The reaction mixture was purified directly by MDAP (HpH) to afford *tert*-butyl (2-(2-(2-(2-(4-(1-(1,3- dimethoxypropan-2-yl)-2-(1,5-dimethyl-6-oxo-1,6-dihydropyridin-3-yl)-1*H*-benzo[d]imidazol-5-yl)piperidin-1-yl)-2-oxoethoxy)ethoxy)ethoxy)ethyl)carbamate (92 mg, 128.88 μmol, 75 %) as a white foam. ^**1**^**H NMR** (400 MHz, DMSO-*d*_6_) δ = 8.01 (1H, d, *J* = 2.20 Hz), 7.69 (1H, d, *J* = 8.56 Hz), 7.65 (1H, dd, J=2.45, 0.98 Hz), 7.46 (1H, d, *J* = 1.22 Hz), 7.10 (1H, dd, *J* = 8.56, 1.71 Hz), 6.68 - 6.76 (1H, m), 4.75 – 4.76 (1H, m), 4.11 - 4.27 (2H, m), 3.95 - 4.02 (3H, m), 3.75 (2H, dd, *J* = 10.27, 4.65 Hz), 3.55 - 3.61 (4H, m), 3.47 - 3.55 (7H, m), 3.37 (2H, t, *J* = 6.11 Hz), 3.16 (6H, s), 3.03 - 3.08 (2H, m), 2.81 - 2.93 (2H, m), 2.62 - 2.71 (2H, m), 2.09 (3H, s), 1.79 - 1.89 (2H, m), 1.60 - 1.73 (1H, m), 1.45 - 1.58 (1H, m), 1.37 (9H, s); **LCMS** (Method A): t_R_ = 0.97 min, [M+H]^+^ 714 (100% purity).

#### 5-(5-(1-(2-(2-(2-(2-Aminoethoxy)ethoxy)ethoxy)acetyl)piperidin-4-yl)-1-(1,3-dimethoxypropan-2-yl)-1H-benzo[d]imidazol-2-yl)-1,3-dimethylpyridin-2(1*H*)-one, 2HCl (29)

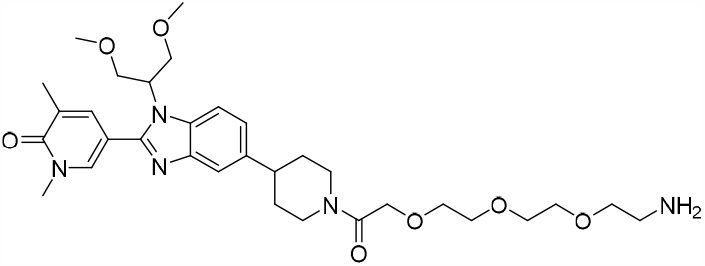

To a stirred solution of tert-butyl (2-(2-(2-(2-(4-(1-(1,3-dimethoxypropan-2-yl)-2-(1,5-dimethyl-6-oxo-1,6-dihydropyridin-3-yl)-1*H*-benzo[d]imidazol-5-yl)piperidin-1-yl)-2-oxoethoxy)ethoxy)ethoxy)ethyl)carbamate (90 mg, 126.07 μmol) in 1,4-dioxane (252 μL) was added 4 M HCl in Dioxane (787.97 μL, 3.1518 mmol) and the reaction mixture was stirred at room temperature for 30 min. The solvent was removed by in vacuo to afford 5-(5-(1-(2-(2-(2-(2-aminoethoxy)ethoxy)ethoxy)acetyl)piperidin-4-yl)-1-(1,3-dimethoxypropan-2-yl)- 1*H*-benzo[d]imidazol-2-yl)-1,3-dimethylpyridin-2(1H)-one, 2HCl (86 mg, 125.24 μmol, 99 %) as a white foam. **LCMS** (Method A): t_R_ = 0.52 min, [M+H]^+^ 614 (97% purity).

#### (3*R*,5*S*)-5-(((*S*)-1-(4-(1-(1,3-Dimethoxypropan-2-yl)-2-(1,5-dimethyl-6-oxo-1,6-dihydropyridin-3-yl)-1*H*-benzo[d]imidazol-5-yl)piperidin-1-yl)-15-(4-(4-methylthiazol-5-yl)phenyl)-1,13-dioxo-3,6,9-trioxa-12-azapentadecan-15-yl)carbamoyl)-1-((*R*)-3-methyl-2-(3-methylisoxazol-5-yl)butanoyl)pyrrolidine-3-sulfonyl fluoride (BRD-SF2)

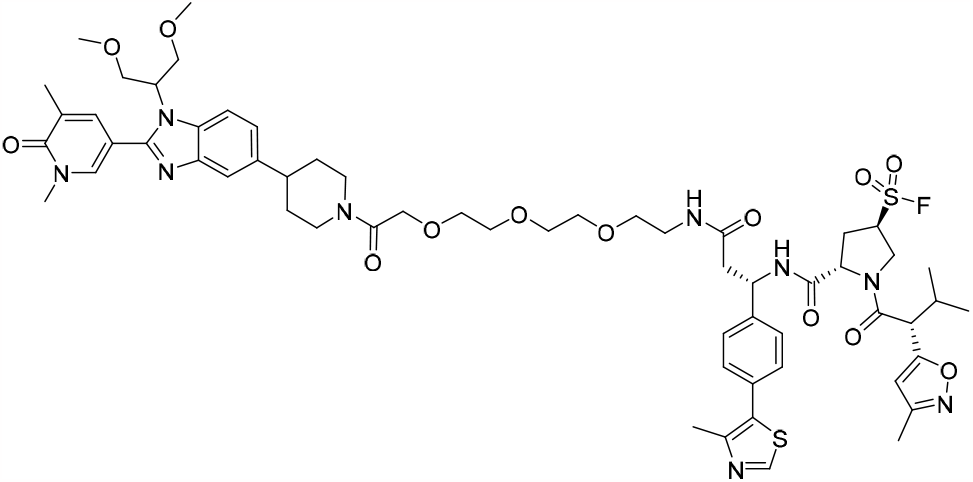

To a stirred solution of (*S*)-3-((2*S*,4*R*)-4-(fluorosulfonyl)-1-((*R*)-3-methyl-2-(3-methylisoxazol-5-yl)butanoyl)pyrrolidine-2-carboxamido)-3-(4-(4-methylthiazol-5-yl)phenyl)propanoic acid, 2TFA (50 mg, 59.9 μmol) and 5-(5-(1-(2-(2-(2-(2-aminoethoxy)ethoxy)ethoxy)acetyl)piperidin-4-yl)-1-(1,3-dimethoxypropan-2-yl)-1*H*-benzo[d]imidazol-2-yl)-1,3-dimethylpyridin-2(1*H*)-one, 2HCl (45.245 mg, 65.89 μmol) in DMF (119.8 μL) and DIPEA (52.2 μL, 299.50 μmol) was added HATU (34.2 mg, 89.85 μmol) and the reaction mixture was stirred at room temperature for 20 min. The reaction mixture was purified directly by MDAP (formic). The volatile solvents were removed and the aqueous was basified to pH 10 with saturated aqueous sodium hydrogen carbonate and extracted with DCM (3 × 50 mL). The organic layers were combined, passed through a hydrophobic frit, and the solvent removed in vacuo to afford (3*R*,5*S*)-5-(((*S*)-1-(4-(1-(1,3-dimethoxypropan-2-yl)-2-(1,5-dimethyl-6-oxo-1,6-dihydropyridin-3-yl)-1*H*-benzo[d]imidazol-5-yl)piperidin-1-yl)-15-(4-(4-methylthiazol-5-yl)phenyl)-1,13-dioxo-3,6,9-trioxa-12-azapentadecan-15-yl)carbamoyl)-1-((*R*)-3-methyl 2- (3-methylisoxazol-5-yl)butanoyl)pyrrolidine-3-sulfonyl fluoride (20 mg, 16.63 μmol, 28%) as a white solid. ^**1**^**H NMR** (400 MHz, DMSO-*d*_6_) δ = 8.93 - 9.02 (1H, m), 8.64 (1H, d, *J* = 8.07 Hz), 8.01 (1H, d, *J* = 2.45 Hz), 7.97 (1H, br t, *J* = 5.75 Hz), 7.87 (1H, br d, *J* = 8.56 Hz), 7.69 (1H, d, *J* = 8.56 Hz), 7.66 (1H, dd, *J* = 2.32, 1.10 Hz), 7.34 - 7.48 (4H, m), 7.10 (1H, dd, *J* = 8.56, 1.47 Hz), 6.21 (1H, s), 5.16 - 5.24 (1H, m), 4.86 - 4.94 (1H, m), 4.82 (1H, m), 4.47 - 4.56 (2H, m), 4.10 - 4.32 (4H, m), 3.89 - 4.03 (4H, m), 3.76 (2H, dd, *J* = 10.27, 4.65 Hz), 3.55 - 3.62 (2H, m), 3.54 (3H, s), 3.47 - 3.51 (2H, m), 3.42 - 3.46 (2H, m), 3.32 (1H, br s), 3.23 - 3.29 (1H, m), 3.16 (6H, s), 3.02 - 3.14 (2H, m), 2.83 - 2.93 (2H, m), 2.65 - 2.74 (2H, m), 2.58 - 2.64 (2H, m), 2.45 - 2.49 (3H, m), 2.23 - 2.35 (2H, m), 2.18 - 2.20 (2H, m), 2.09 (3H, s), 1.81 - 1.89 (2H, m), 1.63 - 1.72 (1H, m), 1.48 - 1.59 (1H, m), 1.22 - 1.31 (2H, m), 0.90 - 1.02 (3H, m), 0.80 (3H, d, *J* = 6.85 Hz); ^**13**^**C NMR** (151 MHz, DMSO-*d*_6_) δ = 168.9, 167.6, 166.9, 161.7, 159.3, 151.8, 151.5, 147.8, 143.2, 142.0, 139.5, 138.9, 137.3, 132.1, 131.0, 130.0, 128.8, 128.7, 127.4 (2C), 127.0 (2C), 126.9, 121.3, 116.7, 112.2, 108.0, 103.5, 103.3, 69.7, 69.7, 69.6, 69.6, 69.1 (2C), 59.7, 58.3 (2C), 48.7, 47.7, 41.7, 40.0, 38.5, 37.5, 33.7, 33.2, 32.4, 31.7, 30.9, 30.0, 20.7, 20.5, 20.2, 19.7, 17.0, 16.0, 16.0, 14.1, 10.9; ^**19**^**F NMR** (376 MHz, DMSO-*d*_6_) δ = 47.79 (1F, s); **LCMS** (Method A): t_R_ = 0.89 min, [(M+2H)/2]^+^601, (95% purity); **HRMS** (C_59_H_77_FN_9_O_13_S_2_) [M+H]^+^ requires 1202.5034, found [M+H]^+^ 1202.5034.

### Synthesis of AR-VHL-SF2 and AR2-VHL-SF2

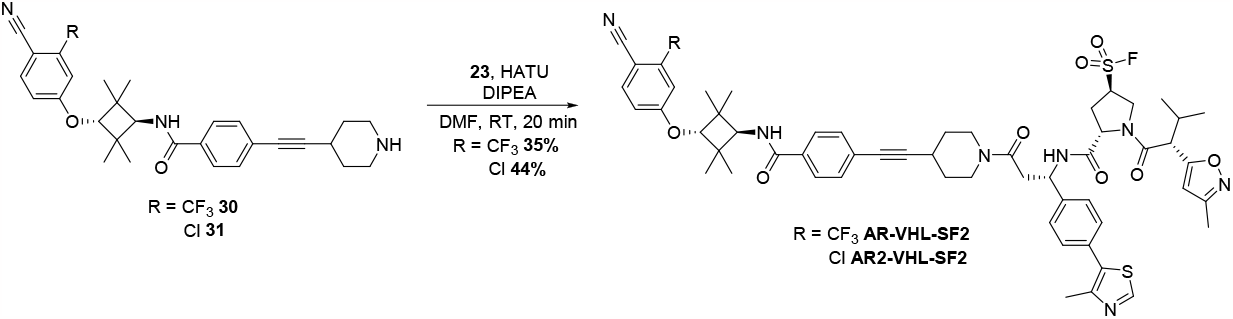

AR binders **30** and **31** were synthesized according to literature precedent.^2^

#### (3*R*,5*S*)-5-(((*S*)-3-(4-((4-(((1*r*,3*r*)-3-(4-Cyano-3-(trifluoromethyl)phenoxy)-2,2,4,4-tetramethylcyclobutyl)carbamoyl)phenyl)ethynyl)piperidin-1-yl)-1-(4-(4-methylthiazol-5-yl)phenyl)-3-oxopropyl)carbamoyl)-1-((*R*)-3-methyl-2-(3-methylisoxazol-5-yl)butanoyl)pyrrolidine-3-sulfonyl fluoride (AR-VHL-SF2)

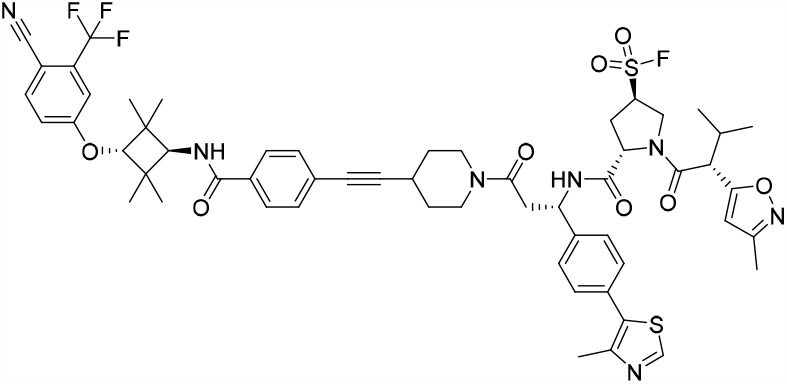

To a stirred solution of (*S*)-3-((2*S*,4*R*)-4-(fluorosulfonyl)-1-((*R*)-3-methyl-2-(3-methylisoxazol-5-yl)butanoyl)pyrrolidine-2-carboxamido)-3-(4-(4-methylthiazol-5-yl)phenyl)propanoic acid, 2TFA (50 mg, 59.9 μmol)and *N*-((1*r*,3*r*)-3-(4-cyano-3-(trifluoromethyl)phenoxy)-2,2,4,4-tetramethylcyclobutyl)-4-(piperidin-4-ylethynyl)benzamide, HCl (33.6 mg, 59.9 μmol) in DMF (119 μL)and DIPEA (52.2 μL, 299.50 μmol) was added HATU (34.2 mg, 89.85 μmol) and the reaction mixture was stirred at room temperature for 20 min. The reaction mixture was purified directly by MDAP (formic). The volatile solvents were removed and the aqueous was neutralised with saturated aqueous sodium hydrogen carbonate and extracted with DCM (3 × 50 mL). The organic layers were combined, passed through a hydrophobic frit and the solvent removed in vacuo to afford (3*R*,5*S*)-5-(((*S*)-3-(4-((4-(((1*r*,3*r*)-3-(4-cyano-3-(trifluoromethyl)phenoxy)-2,2,4,4-tetramethylcyclobutyl)carbamoyl)phenyl)ethynyl)piper-idin-1-yl)-1-(4-(4-methylthiazol-5-yl)phenyl)-3-oxopropyl)carbamoyl)-1-((*R*)-3-methyl-2-(3-methyl-isoxazol-5-yl)butanoyl)pyrrolidine-3-sulfonyl fluoride (23 mg, 20.40 μmol, 35%) as a white solid. ^**1**^**H NMR** (400 MHz, DMSO-*d*_6_) δ = 8.92 - 8.99 (1H, m), 8.60 (1H, br dd, *J* = 7.83, 2.45 Hz), 8.10 (1H, d, *J* = 8.56 Hz), 7.89 (1H, br d, *J* = 8.80 Hz), 7.79 - 7.85 (2H, m), 7.38 - 7.50 (6H, m), 7.30 (1H, dd, *J* = 8.56, 2.45 Hz), 6.18 - 6.23 (1H, m), 5.18 - 5.29 (1H, m), 4.83 - 4.98 (2H, m), 4.52 - 4.57 (1H, m), 4.40 (1H, s), 4.17 - 4.28 (2H, m), 4.09 (1H, d, *J* = 8.80 Hz), 3.92 - 4.00 (1H, m), 3.77 - 3.86 (1H, m), 3.61 - 3.72 (1H, m), 2.84 - 2.98 (3H, m), 2.65 - 2.77 (1H, m), 2.43 - 2.47 (3H, m), 1.73 - 1.88 (3H, m), 1.18 - 1.29 (12H, m), 1.15 (6H, s), 0.95 (4H, m), 0.76 - 0.84 (4H, m); ^**13**^**C NMR** (151 MHz, DMSO-*d*_6_) δ = 174.4, 169.1, 168.4, 166.6, 161.8, 159.3, 159.2, 137.7, 132.8, 132.6, 131.0, 131.0 (d, *J* = 1.66 Hz), 131.0 (2C), 128.8 (2C), 127.8, 127.8 (2C), 127.8, 127.2, 125.6, 123.2, 121.4, 118.7, 115.7, 114.4 (d, *J* = 4.98 Hz), 103.3, 99.6, 84.0, 58.3, 50.1, 48.7, 48.6, 47.7 (d, *J* = 3.87 Hz), 47.1, 41.8, 40.3 (2C), 33.6, 31.3, 30.9 (2C), 30.0, 29.0, 24.5, 24.0 (2C), 23.1 (2C), 20.5, 16.0, 15.9, 13.9, 11.0, 10.9; ^**19**^**F NMR** (376 MHz, DMSO-*d*_6_) δ = (1F, s); **LCMS** (Method A): t_R_ = 1.46 min, [M+H]^+^ 1112, (99% purity); **HRMS** (C_57_H_62_F_4_N_7_O_8_S_2_) [M+H]^+^ requires 1112.4037, found [M+H]^+^ 1112.4027.

#### (3*R*,5*S*)-5-(((*S*)-3-(4-((4-(((1*r*,3*r*)-3-(3-Chloro-4-cyanophenoxy)-2,2,4,4-tetramethylcyclobutyl)- carbamoyl)phenyl)ethynyl)piperidin-1-yl)-1-(4-(4-methylthiazol-5-yl)phenyl)-3-oxopropyl)-carbamoyl)-1- ((*R*)-3-methyl-2-(3-methylisoxazol-5-yl)butanoyl)pyrrolidine-3-sulfonyl fluoride (AR2-VHL-SF2)

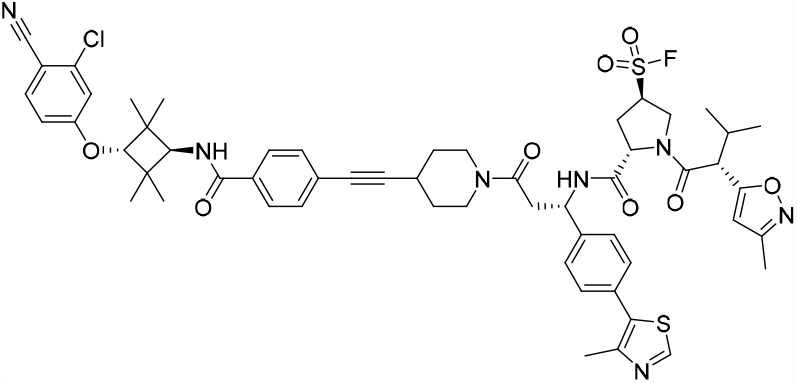

To a stirred solution of (*S*)-3-((2*S*,4*R*)-4-(fluorosulfonyl)-1-((*R*)-3-methyl-2-(3-methylisoxazol-5- yl)butanoyl)pyrrolidine-2-carboxamido)-3-(4-(4-methylthiazol-5-yl)phenyl)propanoic acid, 2TFA (39 mg, 46.72 μmol) and *N*-((1*r*,3*r*)-3-(3-chloro-4-cyanophenoxy)-2,2,4,4-tetramethylcyclobutyl)-4-(piperidin-4-ylethynyl)benzamide, HCl (24.6 mg, 46.72 μmol) in DMF (93 μL)and DIPEA (40.7 μL, 233.61 μmol) was added HATU (26.7 mg, 70.08 μmol) and the reaction mixture was stirred at room temperature for 20 min. The reaction mixture was purified directly by MDAP (formic). The volatiles were removed and the aqueous was basified with saturated aqueous sodium hydrogen carbonate and extracted with DCM (3 × 50 mL). The organic layers were combined, passed through a hydrophobic frit and the solvent removed in vacuo to afford (3*R*,5*S*)-5-(((*S*)-3-(4-((4-(((1*r*,3*r*)-3-(3-chloro-4-cyanophenoxy)-2,2,4,4-tetramethylcyclobutyl)carbamoyl)phenyl)ethynyl)piperidin-1-yl)-1-(4-(4-methylthiazol-5-yl)phenyl)-3-oxopropyl)carbamoyl)-1-((*R*)-3-methyl-2-(3-methylisoxazol-5-yl)butan-oyl)pyrrolidine-3-sulfonyl fluoride (22 mg, 20.40 μmol, 44%) as a white solid. ^**1**^**H NMR** (400 MHz, DMSO-*d*_6_) δ = 8.92 - 8.99 (1H, m), 8.60 (1H, br dd, *J* = 7.82, 3.42 Hz), 7.90 (1H, d, *J* = 8.80 Hz), 7.87 (1H, br d, *J* = 9.29 Hz), 7.77 - 7.84 (2H, m), 7.38 - 7.50 (5H, m), 7.21 (1H, d, *J* = 2.45 Hz), 7.01 (1H, dd, *J* = 8.80, 2.45 Hz), 6.19 - 6.23 (1H, m), 5.16 - 5.29 (1H, m), 4.83 - 4.97 (2H, m), 4.51 - 4.58 (1H, m), 4.32 (1H, s), 4.25 (1H, m), 4.14 - 4.22 (1H, m), 4.07 (1 H, d, *J* = 9.05 Hz), 3.96 (1H, br d, *J* = 9.54 Hz), 3.77 - 3.86 (1H, m), 3.61 - 3.71 (1H, m), 2.85 - 2.98 (3H, m), 2.65 - 2.76 (1H, m), 2.43 - 2.47 (3H, m), 2.24 - 2.35 (2H, m), 2.19 (3H, d, *J* = 3.42 Hz), 1.73 - 1.86 (2H, m), 1.35 - 1.58 (3H, m), 1.23 (6H, s), 1.14 (6H, s), 0.92 - 1.00 (3H, m), 0.76 - 0.84 (4H, m); ^**13**^**C NMR** (101 MHz, DMSO-*d*_6_) δ = 169.2, 169.1, 169.1, 169.0, 166.5, 162.5, 159.3, 151.5, 147.8, 136.8 (2C), 136.0, 133.8, 131.0, 131.0 (2C), 128.7 (2C), 127.8 (2C), 127.1, 125.6 (2C), 116.8, 116.2, 114.6, 103.6, 103.3, 83.9, 58.4, 48.7, 48.5, 40.3 (2C), 30.9 (2C), 30.0 (2C), 23.9 (2C), 23.1 (2C), 20.5, 20.2, 19.8, 19.7, 15.9, 15.9, 11.0, 10.9; ^**19**^**F NMR** (376 MHz, DMSO-*d*_6_) δ = 47.83 (1F, s); **LCMS** (Method A): t_R_ = 1.14 min, [M+H]^+^ 1078 & 1080, (100% purity); **HRMS** (C_56_H_62_ClFN_7_O_8_S_2_) [M+H]^+^ requires 1078.3774, found [M+H]^+^ 1078.3754.

